# Motility Functions as an Essential Defense in the Bacterial Persister Lifecycle

**DOI:** 10.64898/2025.12.17.694804

**Authors:** Yixiao Xiong, Xiaona Fang, Jin Wang

## Abstract

Bacterial persisters are phenotypically antibiotic-tolerant variants driving recurrent infections. While motility is a key trait for bacterial survival and virulence, its function throughout the persister lifecycle remains poorly understood. Here, we unveil a stage-specific trajectory for motility that underlies sophisticated survival strategies in rifampin-induced *Escherichia coli* persistence. During persister formation, intact motility functions as an immediate behavioral defense that decreases intracellular bactericidal antibiotic concentrations to ensure survival. This initial behavioral defense provides critical time for transcriptional reprogramming in persisters, which, by downregulating the major outer membrane porins (OmpC & OmpF) that facilitate antibiotic influx, supplants the preceding motility-based strategy. Subsequently, in structurally fortified persisters, motility becomes dispensable and gradually diminishes– a process independent of cell bioenergetic profile–yet re-emerges as essential for timely resuscitation upon antibiotic removal. Collectively, these findings establish the significance of motility in bacterial survival, redefine persister defense against antibiotics as a transition from behavioral to architectural strategies, and identify motility as a potential adjuvant therapeutic target.

**Graphical Abstract:** 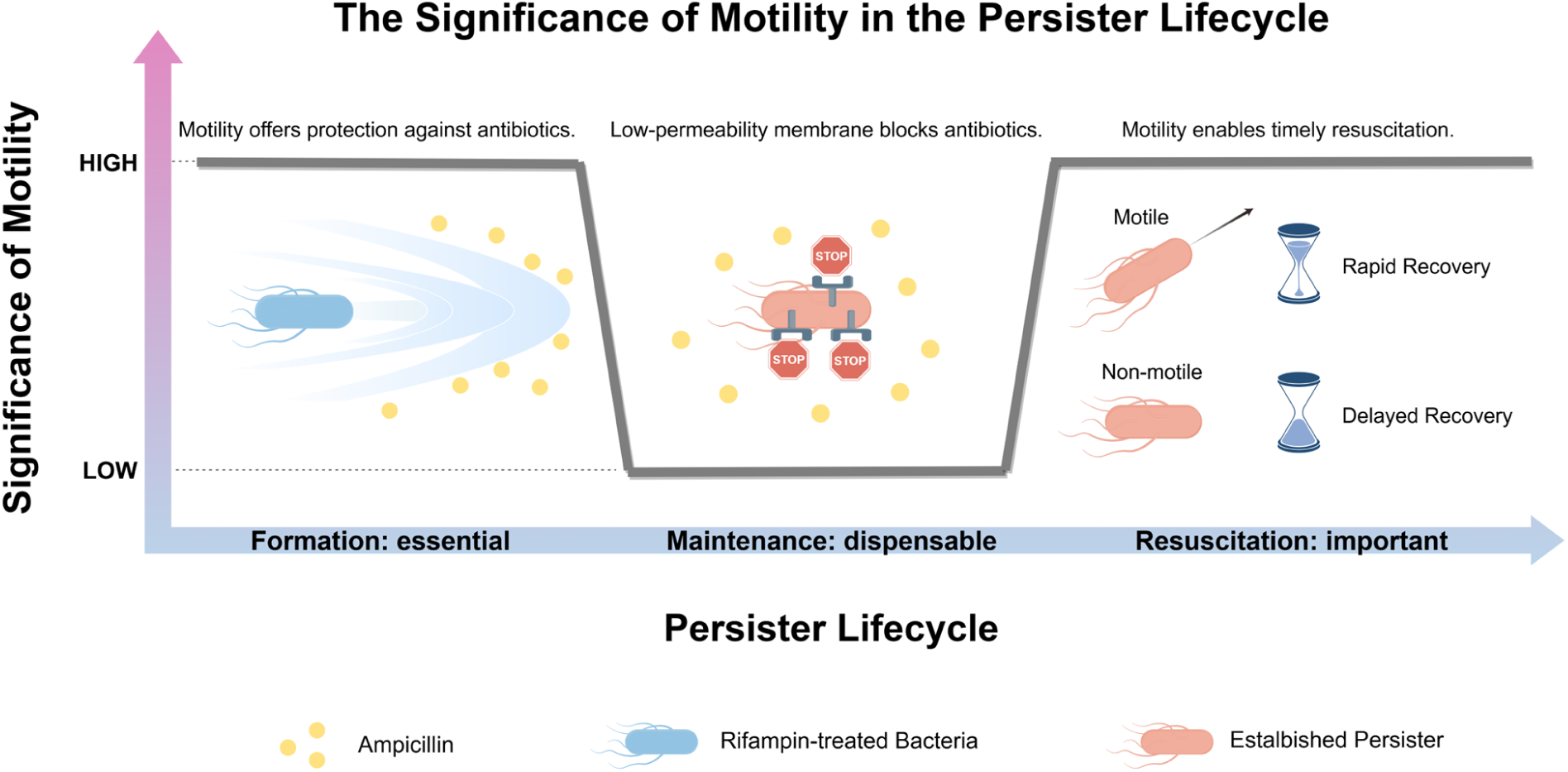

The dynamic roles and underlying mechanisms of bacterial motility throughout the rifampin-induced persister lifecycle: motility initially serves as a behavioral defense mechanism essential for persister formation, then becoming dispensable in the persister state which implements a structural defense via membrane porin downregulation, and ultimately re-emerges as a critical requirement for timely resuscitation.

## Introduction

Bacterial persister cells, often referred to as ‘persisters’, are a subpopulation of cells exhibiting a phenotype that enables them to survive bactericidal antibiotic treatments.^1^ Persisters cannot replicate in the presence of lethal antibiotics but recover and resume growth upon re-culture in a fresh medium without the antibiotic. The re-cultured persister cells are mostly susceptible to antibiotics, while a subpopulation exhibits the persistent phenotype once again.^2,3^ Persisters are extremely difficult to eliminate, thus becoming one of the major causes of antibiotic treatment failure and recalcitrant of bacterial infections.^4^ In clinical practice, multiple chronic bacterial infection diseases are related to persisters, such as tuberculosis,^5,6^ urinary tract infection,^7^ and biofilm infections.^8^

Persistence is driven by a combination of stochastic and responsive factors.^9^ In a steady exponentially growing culture, persisters may emerge spontaneously, which is typically interpreted as bet-hedging: an evolutionary strategy to maximize the fitness of an isogenic population in dynamic environments.^10^ In most observed cases, however, persister formation is induced by external conditions. Factors such as cell density, nutrient starvation and transition, oxidative and acid stress, immune cell exposure, and certain drug exposures may all promote the responsive diversification of bacteria,^2,11–13^ quantitatively and qualitatively modulating the rate of phenotypic conversion into persister cells.^9^ Accordingly, by controlling such conditions when culturing bacteria, various methods have been developed to significantly increase persister formation,^2,14–18^ facilitating mechanistic research into persistence.

*Escherichia coli* (*E. coli*) is among the most extensively studied organisms in persister research. Several non-specific and specific molecular mechanisms have been implicated in *E. coli* persister formation.^19^ For example, the “Persistence As Stuff Happens” (PASH) theory posits that persistence arises from stochastic changes or glitches in cellular metabolism and replication that globally affect bacterial physiology.^2^ Specialized mechanisms, such as the activation of toxin-antitoxin modules, fluctuations in intracellular ATP levels, and the involvement of particular genes, are also recognized as pathways that contribute to persistence.^2,20,21^ Nevertheless, a definitive mechanism or universal determinant that fully explains the persister lifecycle remains elusive.

Recently, multiple elements related to cell motility, including flagellar genes, proton motive force (PMF), and chemotaxis proteins, have been associated with *E. coli* persistence. *E. coli* cells are surrounded by four to six flagella, constituting the structural basis of motility. Each flagellum consists of a reversible rotary motor embedded in the cytoplasmic and outer membranes, a long helical filament extended outside of the cell body, and a short proximal hook connecting the motor to the filament.^22^ Studies have demonstrated that mutation in several flagellar genes could reduce persistence induced by rifampin, tetracycline, and gentamicin.^23,24^ The flagellar motor harnesses transmembrane PMF to generate torque and power the rotation of flagella.^25^ Disruption of PMF has been shown to facilitate the killing of persisters induced by nutrient-shift or starvation.^26,27^ The torque generated by the flagellar motor is then transmitted through a series of delicate structures to drive the rotation of filament^28^ and propel the cell body into two types of movement: “run” (steady forward motion) or “tumble” (directional change with little net displacement). Depending on the environment, bacteria cells can exhibit different types of motion via running and tumbling. On a semi-solid surface, densely packed cells can collectively swirl and migrate, which is called swarming. In liquid environment, planktonic cells can swim as individuals. Naturally, cells move toward beneficial chemical gradients or away from toxic ones, which is achieved by changing the probability of run and tumble through regulation of the chemotaxis signaling pathway.^29,30^ Notably, bacterial chemotaxis proteins, such as CheZ and RbsB, have been shown to be downregulated in persisters.^31^

Motility is essential for the survival of *E. coli*, influencing critical biological processes such as biofilm formation, quorum sensing, bacterial pathogenesis, and host infection.^32^ Yet, motility’s role in bacterial antibiotic survival has received limited attention. Studies have suggested that swarming is mutagenic and can accelerate the rate of antibiotic resistance evolution,^33–36^ but the relevance of swimming motility, especially in persisters, remains poorly understood. Previous studies indicated mixed findings: specifically, *E. coli* persisters induced through HipA overexpression were reportedly non-motile,^37^ while those induced by rifampin pretreatment maintained the ability to swim in liquid or migrate within an agar motility gel.^38,39^ However, these studies neither comprehensively characterize persister motility nor explore the potential interplay between swimming motility and persistence.

In this study, we focus on rifampin-induced *E. coli* persistence against ampicillin. We have uncovered a previously unappreciated role for bacterial motility in the formation and subsequent lifecycle of persister cells. Through quantitative analysis of motile behavior, we demonstrate that persister formation is accompanied by a dynamic remodeling of motility, characterized by a significant decrease in swimming speed and an increase in directional change rate. By chemically or physically inhibiting cell motility during persister formation, we show that intact motility actively functions as a defense mechanism that facilitates persister formation by modulating intracellular antibiotics uptake. We then uncover a mechanistic switch: the initial motility-based defense eventually gives away to a stable, genetically-encoded defense. Transcriptome and functional analysis reveal that established persisters significantly repress outer membrane porin expression, creating a low-permeability shield. With this new structural defense strategy engaged, motility becomes dispensable for persister survival, which was further corroborated by the gradually fading of motility during persistence. This dispensability, however, is temporary, as motility becomes essential once again for the timely resuscitation of persisters upon the removal of antibiotic stress. Additionally, in an effort to elucidate the unsustainability of persister motility, we found that the transient motility is independent of global bioenergetic landscapes, such as nutrient uptake ability and ATP concentrations. These findings collectively establish bacterial motility as a stage-specific determinant in the persister lifecycle--essential for entry, dispensable for maintenance, and critical for timely exit--offering new insights into the mechanisms of persistence and potential strategies to combat persisters and control chronic bacterial infections.

## Results

### Dynamic remodeling of motility characterizes persister formation

Rifampin, a bacteriostatic antibiotic, inhibits mRNA synthesis and blocks transcription by binding and inactivating RNA polymerase. Exposure to rifampin during the mid-exponential phase significantly enhances *E. coli* persistence against bactericidal antibiotics.^14^ In this study, we used *E. coli* MG1655, a derivative from the archetypal K-12 strain with minimal genetic manipulations. To enrich for persister cells and fully eliminate non-persisters, mid-exponential phase bacterial cultures (optical density at 600 nm, OD 600, approximately 0.8) were exposed to 100 μg/ml rifampin for 30 minutes before treated overnight (o/n) with lethal concentrations of ampicillin (300 μg/ml, more than 35 times higher than MIC for wild-type MG1655)^40^. Subsequently, the survival rates were enumerated by plating serial dilutions for colony forming unit (CFU) counting (see schematic in **Fig. 1A** and **fig. S1)**. The results indicated that rifampin pretreatment increased persistence levels against ampicillin to 20.67% ± 4.00%, marking a more than 2000-fold rise compared to un-pretreated controls (0.01%± 0.01%), as shown in **Fig. 1B**.

**Fig. 1.**
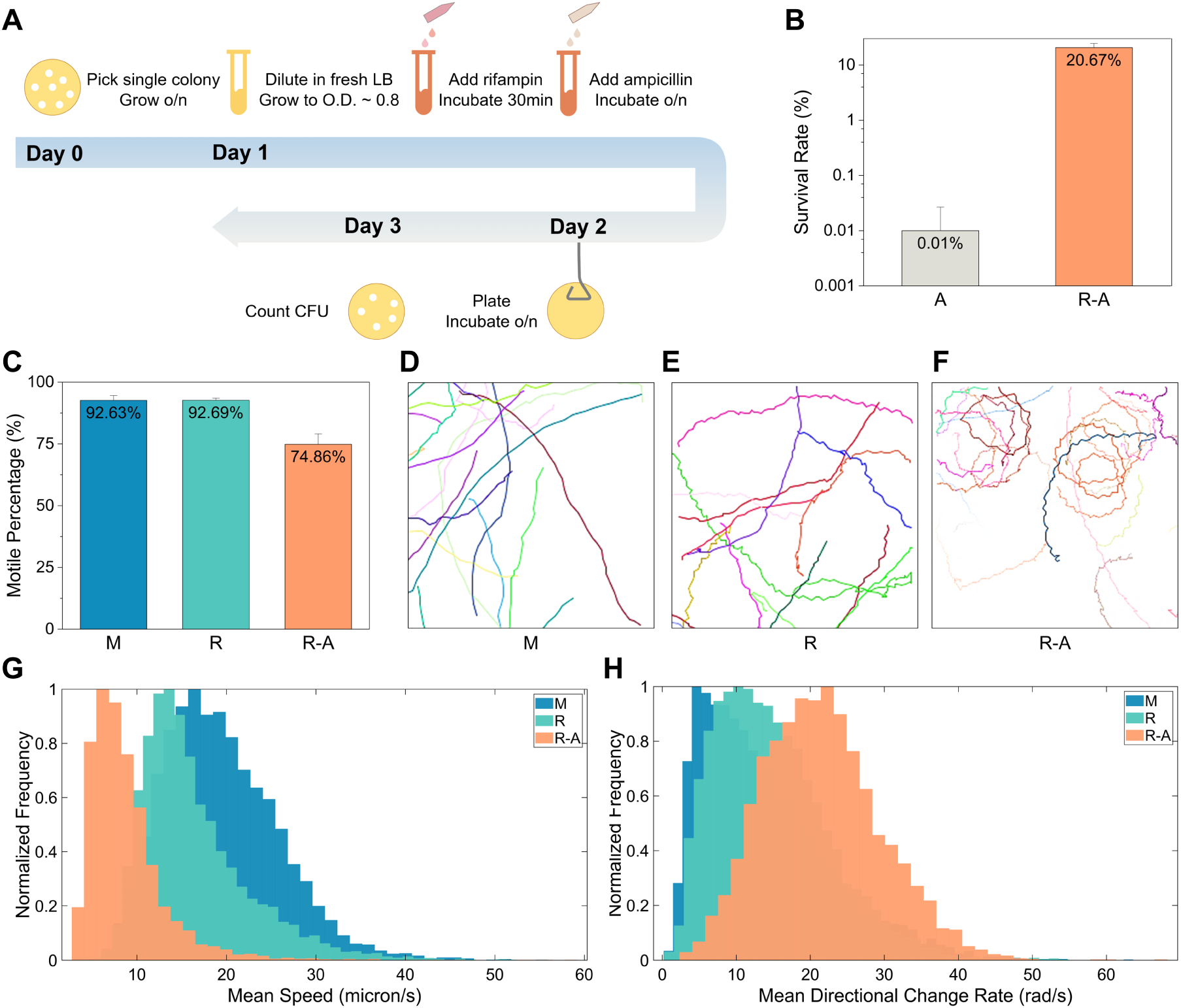
Motility remodeling in rifampin-induced persistence. **A)** Schematic of generating rifampin-induced persisters and evaluating persister survival rates. **B)** Survival rate (mean value ± standard deviation) of *E. coli* MG1655 cells against ampicillin with no pretreatment (*A*) and rifampin-pretreatment (*R-A*). Data represent results from six independent experiments. **C)** Motile percentage (mean value ± standard deviation) of mid-exponential phase cells (*M*), rifampin-treated cells (*R*), rifampin-induced persisters (*R-A*). Data represent results from five experimental videos. The total numbers of cells in the five videos are 598 (*M*), 663 (*R*), and 291 (*R-A*), respectively. **D**-**F)** Representative trajectories of mid-exponential phase cells (**D**), rifampin-treated cells (**E**), and persisters (**F**) swimming in their original living environment (LB or LB with corresponding antibiotics), imaged in the FCS2 chamber. The colored lines show the trajectories of different cells. **G**, **H)** Comparison of track mean speed (**G**) and track mean directional change rate (**H**) distributions for mid-exponential phase cells (*M*), rifampin-treated cells (*R*), and persisters (*R-A*). Data were obtained from at least three experimental videos. Sample sizes for mid-exponential phase cell, rifampin-treated cell, and persister are 6358, 6580, and 4610 tracks, respectively.

A comprehensive elucidation of the motility dynamics in bacterial persister cells is essential to deepen the understanding of their unique physiology and uncover potential mechanistic links between persistence phenotypes and motile behavior. We characterized cell motile behavior across three critical states during rifampin-induced persister formation: the initial mid-exponential growth phase state, the intermediate state following rifampin pretreatment, and the persister state post-overnight ampicillin exposure. To capture the natural swimming behavior of cells in their original living environment, we directly placed 8 μL of bacterial culture onto the Bioptechs FCS2 chamber (0.2 mm plastic pad between slide and coverslip) and used a Nikon ECLIPSE Ti inverted microscope to record bright-field videos at approximately 22 frames per second (FPS). Representative videos of each state--mid-exponential phase, rifampin-pretreated, and persister cells--are provided in **Movie S1 (a**, **b**, and **c)**. The majority of mid-exponential phase and rifampin-pretreated cells were moving rapidly. To our surprise, the persister cells, despite being presumed dormant, also exhibited active motility. Conversely, cells directly exposed to lethal concentration of ampicillin (without rifampin pretreatment) were mostly lysed. As shown in **Movie S1 (d)**, we observed rare appearances of cells with intact morphology but frequently encountered clusters of cell bodies, which made it unlikely to study this population. Focusing on the three critical states, we first quantified the motile percentage as the ratio of motile cells to the total cell population in representative frames drawn from the videos. Moving cells were manually counted, while the total cell count in each frame was quantified by a CellPose model pretrained with our microscopy images.^41,42^ Notably, we considered the cells that could initiate rotation when partly adhered to glass as motile. As shown in **Fig. 1C**, nearly all mid-exponential phase and rifampin-treated cells were motile, with motile percentages reaching 92.63%±1.92% and 92.69%±0.85%. The minority of immotile cells appeared to be fully adhered to the glass surface, resulting in their complete immobilization. Meanwhile, about 74.86%±4.20% of persisters were actively moving.

We then analyzed motility features using the Trackmate plugin of Fiji,^43–45^ where the CellPose model performed cell segmentation, and the Trackmate tracking algorithm linked the cells across successive frames to generate trajectories. Thereupon, the trajectories can provide us with information on the speed and directional change rate of motile cells, quantitatively depicting the two modes of motility in *E. coli*: run and tumble. **Fig. 1D** - **F** shows representative cell trajectories in the initial state, intermediate state, and persister state, respectively. As displayed in **Fig. 1D**, mid-exponential phase cells exhibited diverse swimming trajectories, with some moving generally straight-forward while others had curvature or sudden changes of direction. After rifampin treatment, while the general shapes of the trajectories resembled the mid-exponential phase cells, an appreciable portion became wavy, indicating that the cells kept twisting back and forth (**Fig. 1E**). The trajectories of persisters exhibited apparent distinction from prior stages, as presented in **Fig. 1F**. Nearly all the tracks were quite wavy and curved into circles, displaying a confined motion. This suggested that persister cells moved more difficultly and displayed a higher propensity for tumbling rather than running compared to mid-exponential phase or rifampin-treated cells. Consistent with observed behavioral patterns, cells in the three states exhibited distinct distributions for motility speed and directional change rate. As cells progressed from mid-exponential phase into the persister state, the majority experienced a continuous decline in mean speed and a consecutive increase in mean directional change rate (**Fig. 1G** and **H**). Statistical results of the motility features are listed in **Table 1**. The mean speeds (±standard deviation) for mid-exponential phase, rifampin-treated, and persister cells were 19.4627±6.3734, 16.3823±6.0226, and 8.7784±4.4863 micron/s, respectively. The mean directional change rates (±standard deviation) for these three groups were 13.8226±8.7165, 14.7483±7.6986, and 21.5456±7.9172 rad/s, respectively. Therefore, on the whole, cells gradually swam slower, changed direction more frequently, and exhibited constrained and circular trajectories with larger curvature upon entering persistence. In addition, correlating mean speed with mean directional change rate for each cell across the three states, we found heightened distributions along the inverse diagonal, indicating a negative correlation between the two features. The two-dimensional distributions are shown in **fig. S2**.

**Table 1.** Motility features for different types or states of cell examined in this study. Sample size: mid-exponential phase cells (*M*): 6538 tracks (135825 edges); mid-exponential phase cells in 5% w/v Ficoll (*5%F*): 6689 tracks (207057 edges); mid-exponential phase cells in 10% w/v Ficoll (*10%F*): 5993 tracks (190446 edges); mid-exponential phase cells in 15% w/v Ficoll (*15%F*): 5374 tracks (180475 edges); rifampin-treated cells (*R*): 4771 tracks (118466 edges); persisters (*R-A*): 4610 tracks (263890 edges). Each data (mean ± standard deviation) was obtained from at least 3 videos.

| Sample Name | Track Motility Features (Mean ± SD) |  |
| --- | --- | --- |
|  | Mean speed<br>(micron/s) | Mean directional change rate<br>(rad/s) |
| *M* | 19.4627±6.3734 | 13.8226±8.7165 |
| *5% F* | 11.9863±3.3528 | 15.4470±7.5126 |
| *10% F* | 10.4671±3.2003 | 16.7348±7.2055 |
| *15% F* | 8.2674±3.3862 | 18.8806±7.6203 |
| *CCCP* | ~ 0 | ~ 0 |
| *R* | 16.3823±6.0226 | 14.7483±7.6986 |
| *R-A* | 8.7784±4.4863 | 21.5456±7.9172 |

### Intact motility facilitates the formation of persisters

The observation that cells remain motile—albeit in a remodeled, more constrained manner—during the transition to persistence raises a fundamental question about its function. Is motility merely a residual property of the pre-persister phases, or does the motility itself play an active role in the successful formation of persisters? To investigate the mechanistic relationship between between motility and persister survival while minimizing physiological perturbations, we modulated cell motility through chemical and physical intervention rather than gene knockout strategies and examined how these motility alterations influenced persister fractions during rifampin pretreatment.

We introduced carbonyl cyanide 3-chlorophenylhydrazone (CCCP), a metabolic inhibitor which can depolarize the membrane potential and dissipate PMF, consequently stopping flagellar motor rotation.^46,47^ We first verified the effect of CCCP on bacterial motility through microscopic observation. As shown in **Movie S2 (a)** and **Fig. 2A**, almost all cells in the mid-exponential phase halted motility following exposure to 20 μM CCCP for approximately 2 hours. To assess the effect of impaired motility on persistence levels, cells were treated with 20 μM CCCP at different stages during rifampin-induced persister formation. As illustrated in **fig. S1**, three procedures of adding CCCP were adopted: (1) CCCP was added and incubated for about two hours prior to rifampin treatment and subsequent ampicillin exposure (*C-R-A*). (2) CCCP and rifampin were added concurrently and incubated for 30 minutes before ampicillin exposure (*CR-A*). (3) CCCP was introduced post-rifampin treatment and concurrently with ampicillin (*R-CA*). All bacterial cells treated with ampicillin overnight were plated onto LB agar for CFU counting and survival rate calculating. As shown in **Fig. 2B**, the resulting survival rates all sharply declined to 0.15%∼0.36% – a 100-fold decrease compared to the persisters induced only by rifampin, though still higher than that observed with direct ampicillin treatment. Notably, survival was on the same order of magnitude across the three procedures to add CCCP, showing no significant difference.

**Fig. 2.**
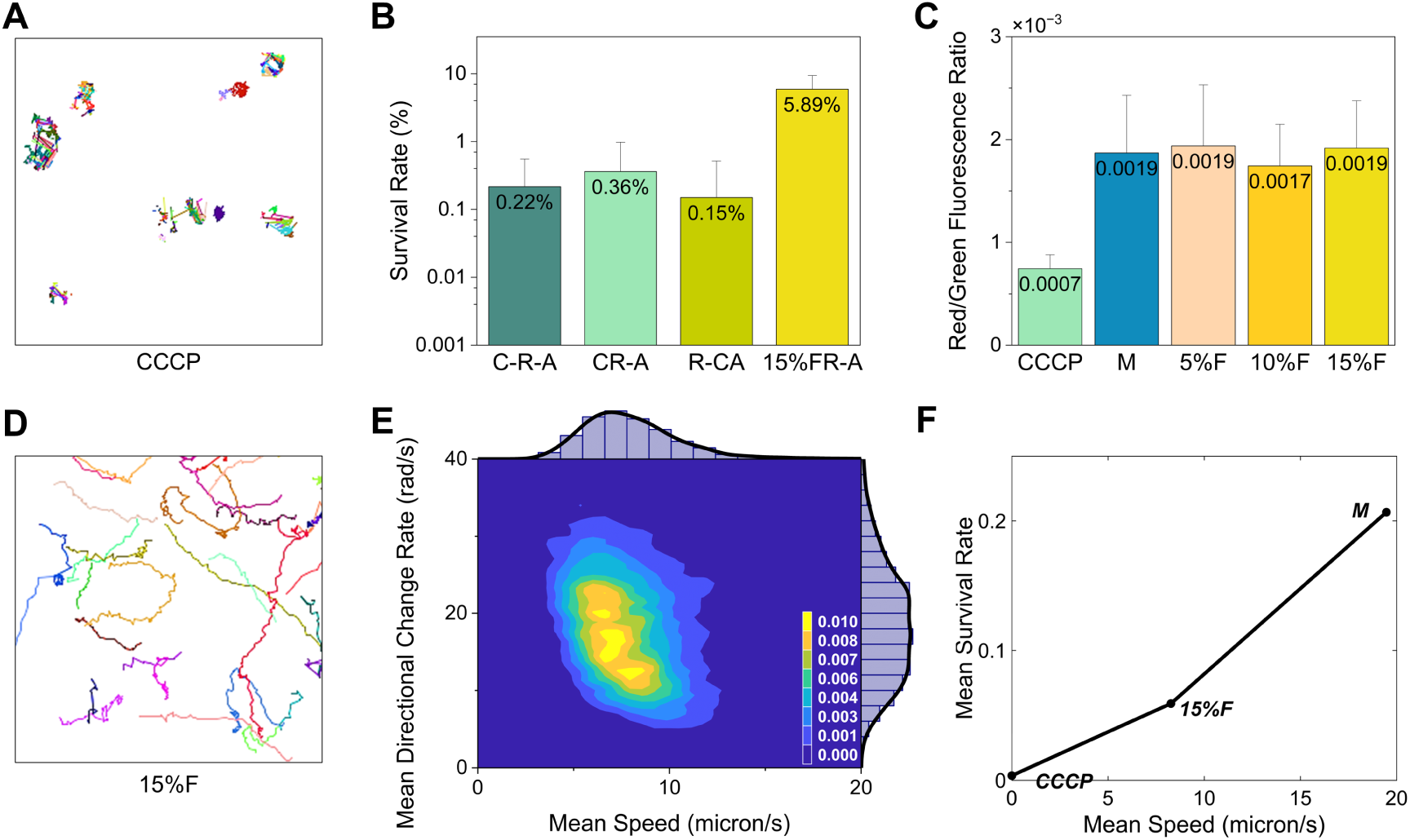
Impact of compromised cell motility on survival rate during persister formation. **A)** Representative trajectories of mid-exponential phase cells swimming in LB after CCCP exposure, observed in the FCS2 chamber. The colored lines show the trajectories of different cells. **B)** Survival rate (mean value ± standard deviation) of *E. coli* cells against ampicillin with exposure to CCCP before rifampin (*C-R-A*), exposure to CCCP and rifampin together (*CR-A*), exposure to CCCP and ampicillin together after rifampin (*R-CA*), and 15% w/v Ficoll in culture medium upon rifampin exposure (*15%FR-A*). Data represent results from six independent experiments. **C)** Membrane potential (mean value ± standard deviation) of mid-exponential phase cells treated with CCCP (*CCCP*), experiencing no treatment (*M*), treated with 5% (*5%F*), 10% (*10%F*), and 15% (*15%F*) w/v Ficoll, as indicated by the red/green median fluorescence intensity ratio measured by flow cytometry. Data represent results from at least three independent experiments. **D)** Representative trajectories of mid-exponential phase cells swimming in LB with 15% w/v Ficoll, imaged in the FCS2 chamber. The colored lines show the trajectories of different cells. **E)** Two-dimensional distribution correlating track mean speed and track mean directional change rate for mid-exponential phase cells in LB with 15% w/v Ficoll, with corresponding marginal distributions. The sample size is 5374 tracks, obtained from seven videos. **F)** Relationship between rifampin-induced survival rate against ampicillin (mean) and swimming speed (mean) for cells under different conditions: *M*, *15%F*, and *CCCP*.

The function of CCCP on ceasing bacterial motility and its role on decreasing rifampin-induced persister fraction indicated the potential importance of cell motility on the formation of persisters. However, prior studies have shown that CCCP pretreatment alone (without rifampin treatment) increased the probability of persister formation.^14^ Our results in **fig. S3** corroborated this finding. Since CCCP exposure stops the rotation of flagellar motor by membrane potential depolarization and PMF disruption, its effects on cells might be beyond motility inhibition. To isolate the impact of motility itself on persistence, we employed Ficoll®400, a viscosity enhancer, to physically slow down cell swimming speed.^48^ First, we verified whether Ficoll could affect cellular physiology, especially membrane potential. Ficoll was introduced into LB bacterial culture at final concentrations of 5%, 10%, and 15% (w/v) during mid-exponential growth phase for 30 minutes (equivalent to the rifampin pretreatment duration, as the two agents were administered concurrently in subsequent experiments). Cell membrane potential was then examined through flow cytometric measurement of a ratiometric fluorescent probe DiOC_2_(3). Membrane potential of mid-exponential phase cells in LB without Ficoll and mid-exponential cells exposed to CCCP were also measured for comparison. As shown in **Fig. 2C**, while CCCP lowered membrane potential to half of that observed in normal cells, the presence of 5%, 10%, 15% w/v Ficoll did not significantly change the membrane potential compared to normal mid-exponential phase cells. This indicated that physically decelerating cell motility did not significantly disturb the membrane potential.

A high concentration of 15% w/v Ficoll was applied to achieve more observable differences in motility and potentially persistent rate. We tracked the motile behaviors of mid-exponential phase cells in LB medium supplemented with 15% w/v Ficoll. Notably, despite the cells maintained their swimming capability in such a highly viscous environment, they appeared struggling, constantly twisting back and forth. The representative trajectories (**Fig. 2D**) displayed wavy paths similar to persister cell trajectories, but less curved generally. Additionally, more cells adhered to the glass compared to those only in LB medium, and some aggregated into clusters (**Movie S2 (b)**). Quantitative analysis confirmed that the presence of Ficoll reduced swimming speed while increasing directional change rate. The average (±standard deviation) of track mean speed and mean directional change rate were 8.2674±3.3862 micron/s and 18.8806±7.6203 rad/s, respectively (also listed in **Table 1**). The correlated two-dimensional distribution of track mean speed and mean directional change rate with marginal distributions of each are presented in **Fig. 2E**. The distribution upon correlation revealed a heightened distribution along the inverse diagonal similar to previous results in **fig. S2**. Subsequently, we co-administered 15% w/v Ficoll and rifampin to mid-exponential phase cells, then measured the survival rates against ampicillin following the same procedure as described previously in this study. The survival rate resulted in 5.89%±3.44%, as shown in **Fig. 2B**, which was higher than those observed with CCCP addition but less than that observed with only rifampin pretreatment.

Taken together, our findings indicated that compromised motility, whether through physical or chemical pathways, diminished rifampin-induced cell survival against ampicillin. This implied that phenotype of cell motility, rather than cellular physiology, played a crucial role on the formation of rifampin-induced persisters. Furthermore, the cells in 15% w/v Ficoll moved more actively than those treated with CCCP (almost completely motionless), and exhibited higher survival rate following rifampin and ampicillin exposure. In contrast, the cells in 15% w/v Ficoll displayed reduced motility compared to those in normal medium, corresponding to a lower survival rate. These observations suggested a correlation between swimming speed and rifampin-induced persister formation probability, as illustrated in **Fig. 2F**, where survival rates increased with swimming speed.

### Motility-dependent modulation of antibiotic uptake underpins persister formation

To investigate how bacterial motility endows rifampin-induced cells with greater chances of surviving bactericidal antibiotic, we evaluated the intracellular uptake of ampicillin in cells with varying motility during persister formation. Fluorescein-labeled antibiotic FITC-ampicillin was used to quantify antibiotic uptake in cells based on fluorescence intensity. The structure and emission spectrum of FITC-ampicillin are displayed in **Fig. 3A**, **B**, while the mass spectrometry is shown in **fig. S4**. To elucidate the relationship between motility and antibiotic uptake, bacterial cultures were supplemented with different concentrations of Ficoll (5%, 10%, and 15% w/v) to systematically modulate cellular motility, generating populations with distinct swimming speeds. We initially characterized the motility of mid-exponential phase cells in response to these experimentally controlled viscosity conditions. As shown in **Movie S2 (b-d)** and **Fig. 3C**, **D**, increasing Ficoll concentration led to a progressive decrease in swimming speed and an increase in directional change rate. Moreover, this observed relationships between motility parameters and Ficoll concentrations were consistent in general, confirming that Ficoll effectively compromised bacterial motility in a dose-dependent manner. Statistical results of motility features are listed in **Table 1**. Additionally, the negative correlation between the two motility parameters displayed in previously described samples (namely the mid-exponential phase cells, rifampin-treated cells, and persisters) was consistent with results in the Ficoll gradient (**fig. S5**). The potential cause of this correlation is discussed in **Note 1**.

**Fig. 3.**
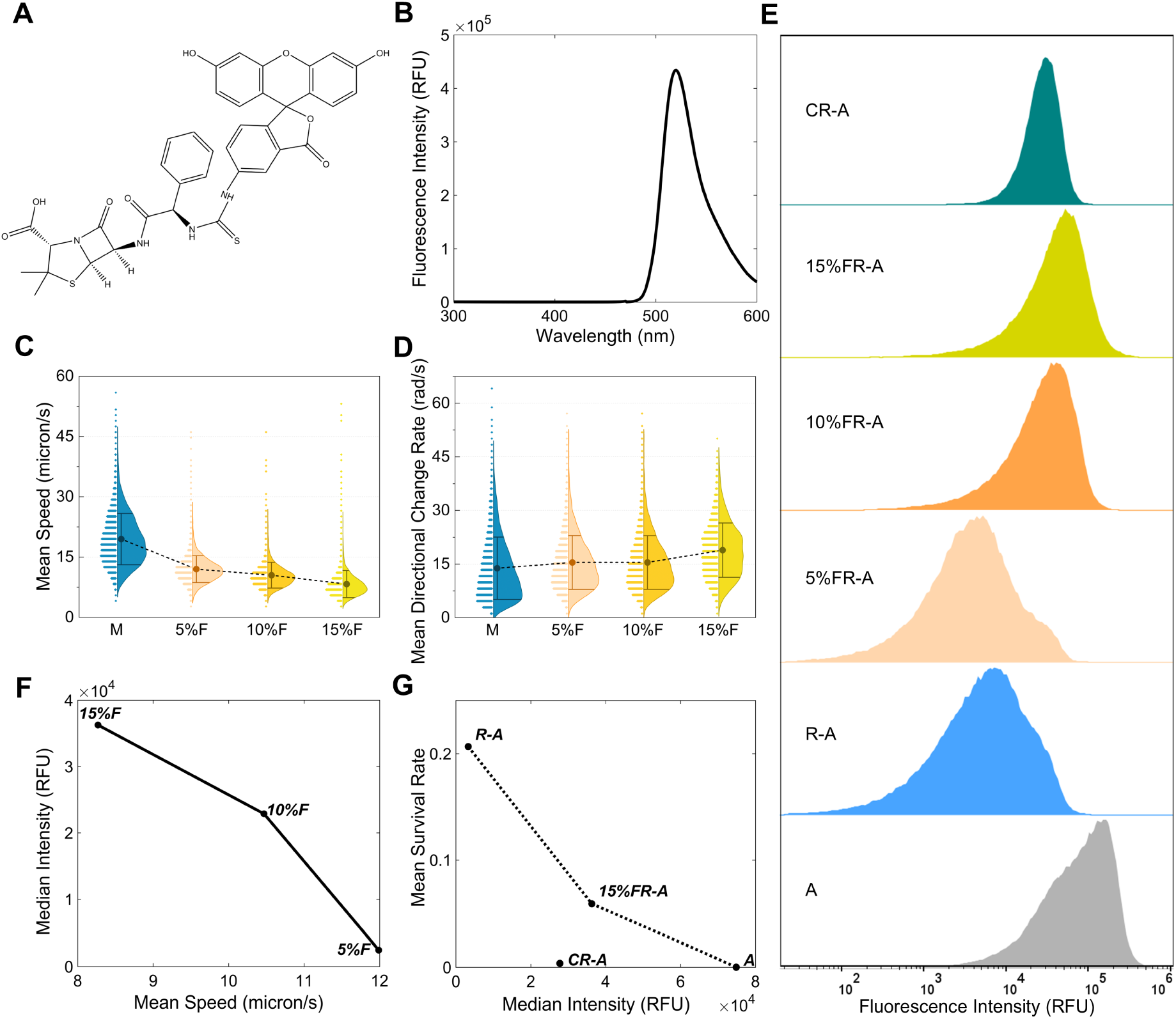
Cell motility influences persister formation by modulating bactericidal antibiotic uptake. **A, B)** Structure (**A**) and emission spectrum (**B**) of FITC-ampicillin. **C**, **D)** Track mean speed (**C**) and track mean directional change rate (**D**) for mid-exponential phase cells (*M*), and mid-exponential phase cells swimming in LB supplemented with 5%, 10%, and 15% w/v Ficoll (*5%F*, *10%F*, and *15%F*). The sample sizes are 6689, 5993, and 5374 tracks, respectively, obtained from at least three videos. The dots represent mean values and the error bars represent ± 1 standard deviation range. **E)** FITC-ampicillin uptake, measured by flow cytometry, in cells after 3 hours of exposure with no pretreatment (*A*, sample size: 149986), exposure to CCCP and rifampin together (*CR-A*, sample size: 193050), rifampin pretreatment in the presence of 15% Ficoll (1*5%FR-A*, sample size: 159685), 10% Ficoll (*10%FR-A*, sample size: 155093), 5% Ficoll (*5%FR-A*, sample size: 142647), and only rifampin pretreatment (*R-A*, sample size: 143513). **F)** Correlation between median FITC fluorescence intensity following 3 hours of exposure and mean swimming speeds of cells experiencing rifampin treatment in the presence of 5%, 10%, 15% w/v Ficoll prior to FITC-ampicillin exposure. **G)** Correlation between survival rate (mean) and antibiotic uptake (median FITC fluorescence intensity following 3 hours of exposure) for the *R-A*, *CR-A*, *15%FR-A*, and *A* experimental groups.

Next, mid-exponential cells experiencing no pretreatment (*A*), rifampin pretreatment (*R-A*), rifampin pretreatment with CCCP addition (*CR-A*), and rifampin pretreatment with 5%, 10%, and 15% w/v Ficoll addition (*5%FR-A*, *10%FR-A*, and *15%FR-A*, respectively) were exposed to 300 μg/ml FITC-ampicillin for 3 hours, respectively (the survival rates of *C-R-A*, *CR-A*, and *R-CA* showed no significant difference, so we chose *CR-A* as the experimental condition). Then the uptakes of ampicillin were determined by flow cytometric measurement of the cellular FITC fluorescence intensity. Representative results are presented in **Fig. 3E**. The *A* cells displayed the highest ampicillin uptake, whereas the cells pretreated with rifampin all exhibited markedly less uptake, confirming rifampin-mediated protection. The *R-A* cells accumulated significantly less ampicillin, with the median intensity value dropping 20-fold compared to the *A* cells. The *CR-A* cells exhibited approximately 10-fold greater uptake than the *R-A* cells, but half that of *A* cells. The ampicillin uptake of the *5%FR-A*, *10%FR-A*, and *15%FR-A* cells fell between the *R-A* and *A* cells, increasing with Ficoll concentrations. Taken into consideration with previous results on motility and survival rate, these findings appeared to suggest a link between cell motility, antibiotic uptake, and the persister survival rates.

To investigate the relationship between cell motility and antibiotic uptake during persister formation, we correlated cell swimming speed with antibiotic accumulation in rifampin-Ficoll-treated cells, as this experimental condition optimally isolated the effects of cell motility. Using the average of mean speed as indicators for motility and the median values of the FITC fluorescence intensity for antibiotic uptake, a clear negative correlation emerged, indicating that slower speeds caused by Ficoll could increase antibiotic uptake during persister formation (**Fig. 3F**). Notably, the antibiotic uptake levels of *CR-A* cells did not exceed those of *15%FR-A* cells, as one would expect due to the complete loss of motility. This deviation may be attributed to CCCP’s broader impact on cellular physiology beyond its effects on cell motility. Nevertheless, these results collectively suggest that motility changes that decreased swimming speed--whether due to the off-target effects by CCCP, or the direct physical effects by Ficoll--can influence cellular antibiotic uptake during persister formation.

Finally, to evaluate the overall contribution of antibiotic uptake to persister survival and elucidate the underlying mechanism, we correlated the antibiotic uptake levels observed under different conditions with the corresponding survival rates determined in **Fig. 1B** and **2B**. As illustrated in **Fig. 3G**, a negative correlation between antibiotic uptake and survival rate was consistent across various experimental conditions except for the *CR-A* group, demonstrating the significant impact of antibiotic uptake on bacterial survival. The observed deviation in the *CR-A* group was consistent with the lower antibiotic uptake as displayed in **Fig. 3E**, likely stemming from CCCP’s additional physiological effects that function independently of antibiotic uptake mechanisms in mediating survival. Overall, these findings collectively confirm that intact motility acts as a critical defense strategy during rifampin-induced persister formation, primarily by modulating the bactericidal antibiotic uptake in cells. Compromised motility can increase intracellular antibiotic levels and render cells vulnerable.

### A mechanistic switch to porin repression marks a motility-dispensable state in persisters

To further investigate the temporal dynamics of motility’s impact on antibiotic uptake, we additionally monitored intracellular antibiotic levels at the onset (0h) and after overnight ampicillin exposure (o/n). As shown in **fig. S6**, flow cytometry revealed that at the initial time point, the *15%FR-A* cells accumulated more antibiotic than *R-A* cells but less than *A* cells, consistent with observations at 3h ampicillin exposure. This indicated that the motility-mediated barrier to antibiotic uptake was established immediately upon ampicillin exposure. Reduced motility increased antibiotic uptake from the onset of exposure, further reinforcing its role during the early stage of persister formation. However, after overnight ampicillin treatment, the intracellular antibiotic levels declined in both *15%FR-A* and *R-A* populations and the difference between the two groups diminished, whereas the *A* population maintained high levels of uptake. These temporal dynamics potentially suggest that motility-driven differences in antibiotic uptake are most prominent during early stage of persister formation, but may play a diminishing role once persister cells are fully established.

To unveil the molecular mechanisms responsible for this stable, low-uptake persister state, we performed transcriptome profiling (RNA sequencing) of established persisters (*P*) and compared them to mid-exponential phase (*M*) and rifampin-treated (*R*) populations. We focused our analysis on a set of genes known to be involved in ampicillin transport and enzymatic degradation, whose pathways are summarized from existing literature in **Fig. 4A**.

**Fig. 4.**
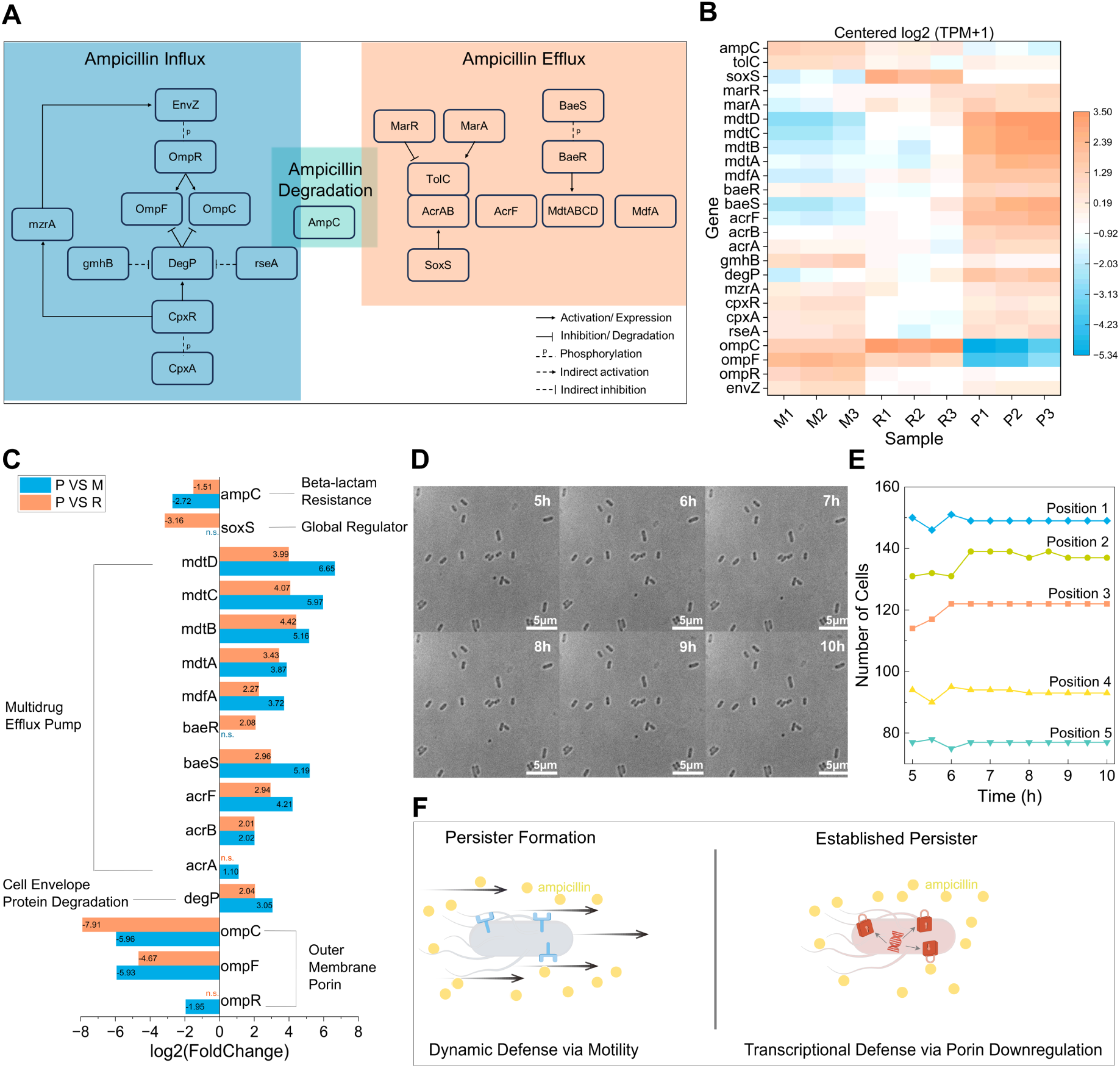
Transcriptional defense via OMP downregulation and the dispensability of motility in established persisters. **A)** Schematic diagram of key gene interaction pathways involved in ampicillin influx, efflux and enzymatic degradation in *E. coli*. **B)** Gene expression heatmap for the relevant genes in mid-exponential phase cells (*M*), rifampin-treated cells (*R*), and persisters (*P*). Three biological replicates were included for each population. Expression shown as log_2_(TPM+1). **C)** Bar plot showing the log_2_(fold change) of genes significantly differentially expressed (|log2(FoldChange)| > 1, qValue < 0.05) in persisters (*P*) compared to mid-exponential phase cells (*M*) and rifampin-treated cells (*R*), with functional annotations. The ‘n.s.’ indicates that the expression difference is not significant. **D)** Real-time microscopy images of a representative field of view on the agarose gel pad, from five to ten hours after persister CCCP exposure. **E)** Quantification of cell counts over time in five fields of view following persister CCCP treatment and in ampicillin flow. **F)** Conceptual model illustrating the mechanism switch from the motility-dependent dynamic defense during persister formation to the porin-downregulation-based transcriptional defense in established persisters.

The primary route of ampicillin entry is through outer membrane porins (OMPs), especially OmpF and OmpC,^49,50^ whose expression is positively regulated by the EnvZ-OmpR two-component system^51^ and MzrA-EnvZ interactions.^52,53^ Conversely, while DegP can serve as a chaperon for correctly folded OMPs,^54^ this periplasmic protease can also degrade OmpF and OmpC, thus negatively controlling porin abundance and reducing influx.^55^ DegP expression is positively regulated by the two-component system CpxA-CpxR (directly)^56,57^ and mutations in *gmhB* or *rseA* (indirectly, mutations in gmhB and rseA increase DegP expression through sigma-E regulon activation).^55^ To counteract antibiotic entry, *E. coli* employs multidrug efflux pumps to expel harmful substances. The prominent AcrAB-TolC system can actively export ampicillin,^58^ which is activated by MarA (MarA counteracts MarR-mediated repression) and oxidative stress sensor SoxS.^59,60^ Additionally, some efflux pumps in *E. coli* have not been experimentally verified to contribute to ampicillin export but may potentially be involved, were included in our analysis as well. For example, AcrF is a homolog of AcrB and can export some β-lactams.^59^ MdtABCD, activated by the BaeS-BaeR two-component system,^61^ and MdfA,^62^ can all export a wide variety of drugs. Finally, beyond these transport mechanisms, direct enzymatic inactivation of the antibiotic can be mediated by the periplasmic β-lactamase, AmpC.^49^

A global view of the expression dynamics, presented as a heatmap of log_2_(TPM+1) values for *M*, *R*, and *P* population (3 samples each), reveals that established persisters possessed a transcriptional profile strikingly different from both *M* and *R* cells (**Fig. 4B**). The most apparent changes pointed to the genes controlling the OMPs, which were significantly down regulated, and efflux pumps, which were mostly up-regulated.

Analysis of significantly differentially expressed genes confirmed this transcriptional overhaul and identified its key drivers (**Fig. 4C**). The most profound change was the severe downregulation of genes encoding the major OMPs, with the intracellular RNA abundance of *ompC* and *ompF* dropped by approximately 100-fold in persisters. Specifically, the log_2_(FoldChange) values for *ompF* was *P* vs. *M* = -5.93; *P* vs. *R* = -4.67 and for *ompC* was *P* vs. *M* = -5.96; *P* vs. *R* = -7.91. The down regulation was significant not only relative to *M* cells, but, critically, also to *R* cells. This timing indicated that the profound loss of OMPs was a hallmark of the fully established persister state, rather than an intermediate effect of rifampin pretreatment. The OMP repression was further supported by the downregulation of their upstream regulator *ompR* and the significant upregulation of *degP* with log2(FoldChange) values of *P* vs. *M* = 3.05 and *P* vs. *R* = 2.04. Concurrently, we observed upregulation of multiple multidrug efflux pump systems, including components of the AcrAB-TolC system (*acrA* and *acrB*), the BaeSR two component system (*baeS* and *baeR*), and the Mdt family of transporters (*mdtA*, *mdtB*, *mdtC*, *mdtD*), as well as the *acrF* and *mdfA* efflux pump genes. The upregulation of efflux-related genes in persisters has been revealed by previous research.^63^ However, while these efflux genes were broadly upregulated, their induction was less pronounced when comparing *P* cells directly to *R* cells in our system. In contrast, the OMP genes underwent their most severe repression during this final *R* to *P* transition. This highlighted that the dominant mechanism fortifying the established persisters was the comprehensive shutdown of antibiotic influx pathways, rather than a comparatively modest enhancement of efflux. Furthermore, the functional impact of the efflux pump upregulation may be constrained, as the outer membrane channel gene *tolC*, required for the function of many of these pumps,^61,64,65^ was not significantly upregulated. Finally, the gene encoding the β-lactamase *ampC* was downregulated, suggesting that established persisters do not rely on enzymatic degradation of the antibiotic.

These transcriptional data pointed to the emergence of a static defense mechanism that primarily reduced membrane permeability, rather than enhancing efflux activity, to protect the established persisters against antibiotics. A critical question, however, is whether this new strategy replaces the dynamic, motility-based defense or merely supplements it. In other words, is motility still essential for survival once the persister cells’ membrane is transcriptional fortified?

To address this, we directly assessed the necessity of motility for the survival of established persisters. 20 μM CCCP was introduced to abolish motility in persister cells (confirmed by microscopy, shown in **Movie S3**). Subsequently, the immobilized persister cells were fixed on an LB agarose gel pad containing 300 μg/ml ampicillin in the FCS2 chamber. LB medium supplemented with an identical ampicillin concentration (300 μg/ml) was continuously flowed through the chamber during the experiment. Cell development was monitored via real-time microscopy at 30 distinct positions over 10 hours, with images captured every 30 minutes. As shown in a representative time-lapse montage (**Fig. 4D**) and cell counts from five representative positions (**Fig. 4E**), no significant cell lysis or growth was observed. The cell counts remained stable over time, with only slight variations due to sample drift. These results demonstrate that even when persister motility is chemically (CCCP exposure) and physically (fixed on agarose gel pad) inhibited, persister cells remained intact, indicating that active movement is not essential for their survival under antibiotic stress.

Collectively, our results support a model of mechanism switching, as illustrated in **Fig. 4F**. During the initial, dynamic stage of persister formation, undisrupted motility provided protection against the antibiotic. Subsequently, cells transitioned to a stable, genetically-encoded defense dominated by the severe transcriptional repression of major OMPs. Our findings extend prior work by providing new evidence that downregulation of OMPs is associated with persistence.^63^ Under this “gate-locking” strategy, motility that provided the initial protection became dispensable. This confirmation of a switch from a dynamic to a static defense raised new questions about the subsequent persister lifecycle: what is the long-term fate of this now non-essential motility, and what role, if any, does it play in the eventual resuscitation?

### Motility fades during persistence but is essential for timely resuscitation

To determine the fate of motility in the subsequent stages of the persister lifecycle, we first sought to characterize its long-term dynamics during persistence. We approached this question by prolonging the observation of persister cells in their original culture (LB with antibiotics) to determine the undisturbed, pristine fate of motility over time. Cell samples were collected at 0h, 6h, 12h, 21h, and 34h after overnight ampicillin exposure (**Movie S4**), examined via microscopy, and quantified for total cell counts in field of view and motile percentage. As shown in **Fig. 5A**, while the average cell count fluctuated without statistically significant difference over time (indicated by the Tukey paired comparison results in **Fig. 5B**), the average percentage of motile cells continuously declined from over 70% to nearly 0%. These results indicated that although persister cells did not undergo mass lysis or death during extended ampicillin exposure, they did not maintain active motility indefinitely, consistent with its non-essential role for survival in this state. Notably, a fraction of the persister cells exhibited elongation over time without division, likely due to the decreased efficiency of ampicillin following long periods of incubation at 37°C.

**Fig. 5.**
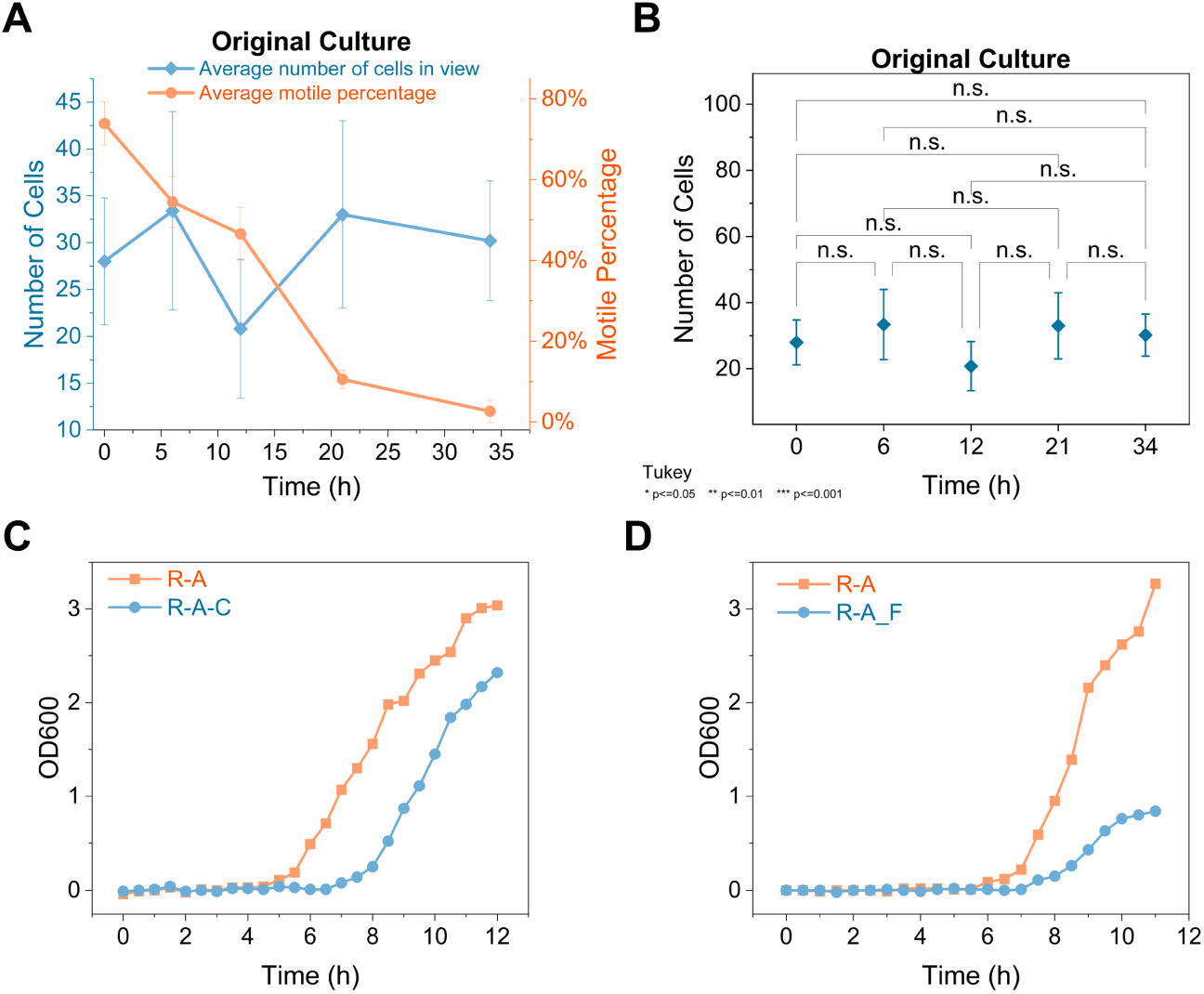
Motility gradually diminishes during persistence but is essential for timely resuscitation. **A)** Cell count (blue, left axis) and motile percentage (orange, right axis) of persisters in their original culture at 0h, 6h, 12h, 21h, 34h after overnight ampicillin exposure (approximately 16h, 22h, 28h, 37h, 50h after rifampin-pretreatment). Data at each time point represent the mean (±standard deviation) from at least three videos. **B)** Tukey paired comparison results of number of persister cells in their original culture after 0h, 6h, 12h, 21h, 34h. At the 0.05 level, Shapiro-Wilk normality test suggests that the data at each time point was significantly drawn from a normally distributed population. The abbreviation ‘n.s.’ indicates no statistically significant difference between the samples in a test (all P values > 0.05). **C)** Representative growth curves of persister cells (*R-A*) and persister cells exposed to CCCP (*R-A-C*) during resuscitation in fresh medium. **D)** Representative growth curves of persister cells during resuscitation in fresh medium with (*R-A_F*) or without (*R-A*) 15% w/v Ficoll.

The original culture medium, initially introduced during mid-exponential phase prior to rifampin pretreatment, had been used for over 50 hours by the end of the observation period. The extended duration of medium use introduced complications in identifying causes of persister motility loss, including nutrient depletion in the culture medium and the accumulation of secondary metabolites. We addressed this by confirming that persisters showed the same decay in motility when transferred to fresh medium (containing antibiotics) or PBS (could not support metabolic activity) (see **Note 2**).

Having established that motility naturally declined during persister maintenance, we next investigated its role in the resuscitation of persister cell upon removal of the antibiotic. Persister cells were treated with CCCP to stop motility and then diluted into fresh, antibiotic-free LB medium to initiate resuscitation. Growth status, monitored via OD _600_ measurement, was compared between CCCP-treated and untreated persister cells during resuscitation. As displayed in **Fig. 5C**, we observed a significant delay in the resumption of growth within the CCCP-treated persister population compared to the untreated group. This observation was further corroborated by independent experiments (**fig. S7a** and **b**). Though the precise timing of resuscitation varied among experiments, the delay was consistent, ranging from one to two hours. Notably, the immobilized persisters that were exposed to CCCP regained motility after resuscitation. These results suggest that motility may play a critical role in facilitating timely exit from the persister state.

To further validate the role of motility in persister resuscitation and exclude potential confounding effects of CCCP, we also assessed the growth resumption of persister cells in the presence of 15% w/v Ficoll. Persister cells were diluted into fresh LB medium containing 15% w/v Ficoll, and their growth was monitored. As a result, the presence of Ficoll delayed the resuscitation of persister cells compared to the control group in normal medium (**Fig. 5D**). Independent experiments showed consistent delays of about one hour in growth resumption (**fig. S7c** and **d**). The observed delay is most likely attributable to the increased viscosity hindering bacterial movement, verifying the essentiality of motility during persister resuscitation.

Therefore, our findings highlight a stage-specific requirement for cell motility in the persister lifecycle. Initially, during the persisters formation, cell motility serves as a critical, instant defense, preventing antibiotics from entering cells. This behavior defense affords the bacterial cells sufficient time to establish a more permanent architectural defense by transcriptionally downregulating the membrane porins responsible for antibiotic influx. In this structurally fortified state, motility becomes dispensable for persisters maintenance. However, motility re-emerges as essential for the timely resuscitation upon antibiotic removal. These observations indicate that motility’s role is not static but follows a dynamic ‘high-low-high’ pattern of significance throughout the persister lifecycle, providing crucial advantages at different stages of survival and recovery.

### Persister motility is independent of global bioenergetic landscapes

Previous studies have revealed that the assembly and operation costs of the flagella is estimated to total approximately 10% of the entire energy budget of an *E. coli* cell,^33,66^ implying that the capability of nutrient uptake and ATP synthesis might have potential impact on cell motility. We therefore investigated whether the unsustainable motility in persister cells could stem from internal physiological limitations, such as impaired nutrient uptake or energy utilization. To figure out the energy profile of persister cells, we first examined the glucose uptake ability of cells through flow cytometric measurement of a fluorescent D-glucose analog 2-(N-(7-nitrobenz-2-oxa-1,3-diazol-4-yl) amino)-2-deoxy-d-glucose (2-NBDG).^67^ As illustrated in **Fig. 6A**, rifampin-induced persisters (*R-A*) exhibited robust glucose uptake ability comparable to mid-exponential phase cells (*M*) and rifampin-treated cells (*R*). In contrast, the cells directly treated with ampicillin (*A*) displayed significantly diminished glucose uptake. Notably, prolonged culture in original medium for another 24 hours (*R-A_24h*) or 31 hours (*R-A_31h*) following overnight ampicillin exposure resulted in observable enhancement of glucose uptake. These findings indicate that persister cells maintained nutrient uptake capacity for a long period of time, thereby ruling out the possibility that motility cessation was due to impaired nutrient uptake. There remains the possibility that persisters might not be able to utilize intracellular energy, for instance, ATP, to indirectly power motility. To verify this, we engineered a reporter system by fusing the rrnB P1 promoter with a fast-folding and fast-degrading GFP variant (superfolder-GFP-ssrA, sfGFPssrA) to monitor intracellular ATP levels. The 16S rRNA rrnB P1 has been established as an ATP-sensing promoter in bacteria, with its transcription levels correlating positively with ATP concentration.^21,68,69^ The protease degradation tag ssrA was fused to the C-terminal of superfolder-GFP to accelerate protein turnover, thereby enabling rapid detection of promoter activation through increased fluorescence and deactivation through decreased signal.^70^ Flow cytometric measurements (**Fig. 6B**) revealed a marked increase in intracellular ATP concentration in rifampin-treated cells (*R*) relative to mid-exponential phase cells (*M*), whereas ampicillin exposure (*A*) triggered a depletion of intracellular ATP pools. Notably, rifampin-induced persisters (*R-A*) displayed bimodal ATP distribution: one subpopulation exhibiting elevated ATP levels similar to *R* cells, while the other displayed reduced ATP levels comparable to *A*cells. This intriguing finding suggested that sequential exposure to rifampin and ampicillin could generate persisters with heterogeneous ATP states which combined the intracellular energy characteristic of separate exposure to each antibiotic.

**Fig. 6.**
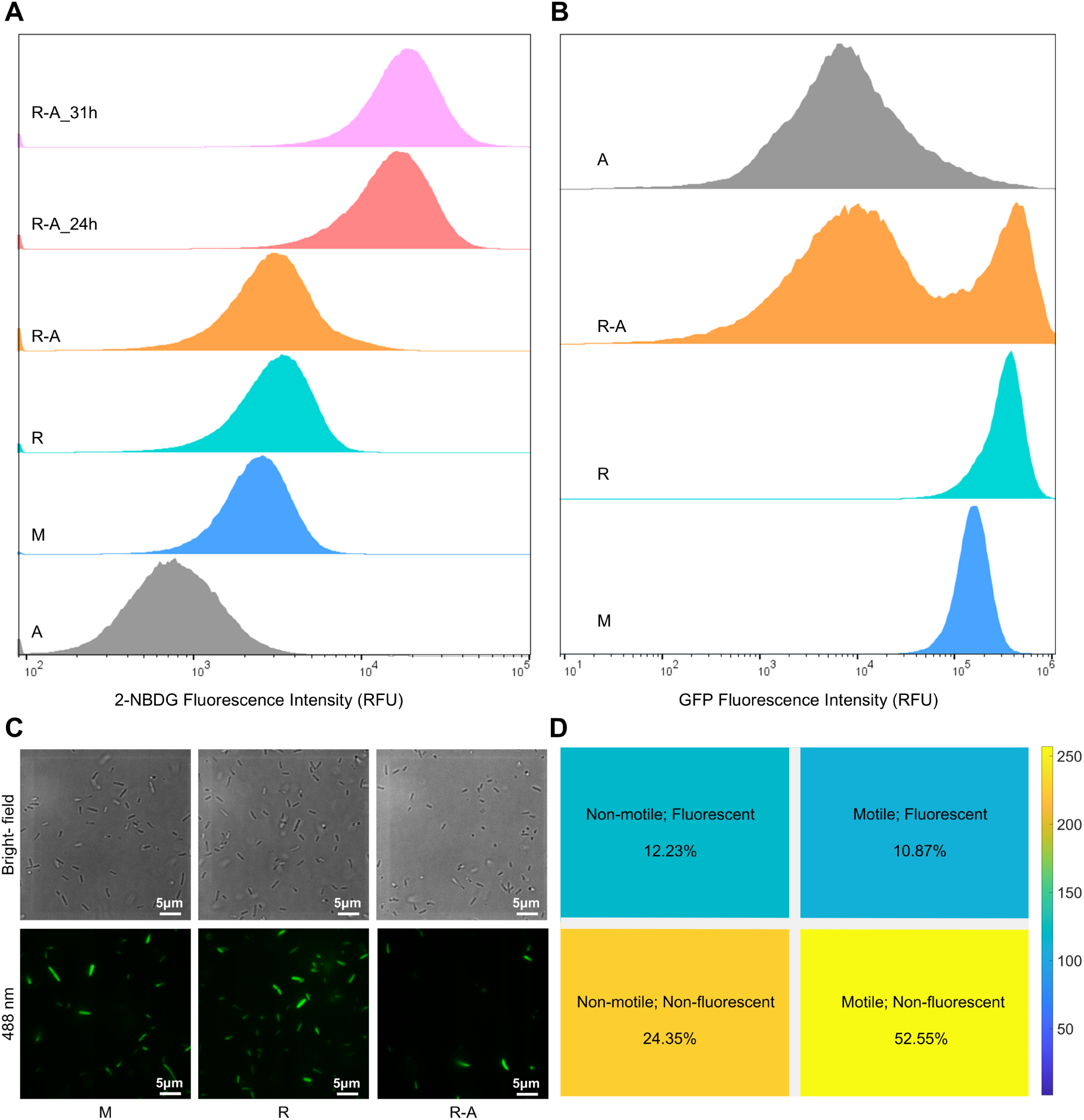
Persister energy profile and motility. **A)** Flow cytometry results of glucose uptake levels (2-NBDG fluorescence intensity) for (from bottom to top) ampicillin exposed cells (*A*, sample size: 126724 cells), mid-exponential phase cells (*M*, sample size: 186755 cells), rifampin treated cells (*R*, sample size: 190510 cells), rifampin-induced persisters after overnight ampicillin exposure (*R-A*, sample size: 194433 cells), rifampin-induced persisters 24 hours after overnight ampicillin exposure (*R-A_24h*, sample size: 189999 cells), and rifampin-induced persisters 31 hours after overnight ampicillin exposure (*R-A_31h*, sample size: 213168 cells). **B)** Flow cytometry results of intracellular ATP levels (indicated by GFP fluorescence intensity) for (from bottom to top) mid-exponential phase cells (*M*, sample size: 89942 cells), rifampin treated cells (*R*, sample size: 97529 cells), rifampin-induced persisters (R-A, sample size: 64730 cells), and ampicillin exposed cells (*A*, sample size: 81464 cells). **C)** Bright-field and fluorescence microscopy images of cells with the ATP reporter at mid-exponential phase, after rifampin-pretreatment, and during rifampin-induced persistence, respectively. The bright-field and fluorescence images were drawn from separate motility videos on the same position of the same sample, so the cells in brightfield and fluorescence image might not match in position due to cell movement. Nearly all mid-exponential phase and rifampin-treated cells in view exhibited high fluorescence, as indicated by the close cell counts in the bright-field and fluorescence microscopy images. In contrast, only a portion of the persisters exhibited fluorescence, evident from the noticeably fewer bright cells in the fluorescence image than in the corresponding bright-field view. **D)** Frequency distribution correlating intracellular ATP levels (indicated by fluorescence intensity) and motility for persisters. Proportions of different states are shown. The sample size is 982 cells from 16 videos across three independent experiments.

The heterogeneity of intracellular ATP levels in rifampin-induced persister cells prompted another critical question: given that some persisters exhibited high ATP levels, could this energy be exploited for motility? In other words, is there a correlation between ATP abundance and cell motility in persisters? To verify this, we monitored cell motility and ATP levels (using GFP fluorescence) of mid-exponential phase cells, rifampin-treated cells, and rifampin-induced persisters, under brightfield and fluorescence microscopy (representative frames displayed in **Fig. 6C**). Nearly all the mid-exponential phase and rifampin-treated cells were motile (as depicted in **Fig. 1C**) and displayed bright fluorescence (indicated by the similar cell count in view under bright-field and fluorescence microscopy images in **Fig. 6C**, left and middle column). This observation was consistent with the flow cytometry results in **Fig. 6B**, implying that high levels of ATP seem to indicate active cell motility. However, among the motility-heterogeneous persisters, only a fraction exhibited bright fluorescence, as indicated by the clear difference between the cell counts under bright-field and fluorescence microscopy images (**Fig. 6C**, right column). Despite significant heterogeneity in both ATP levels and cell motility within the persisters, over 74% of cells exhibited motility (**Fig. 1C**) while only about 26.7% of cells maintained high ATP levels (**Fig. 6B** and **fig. S8**), indicating a lack of correlation between these two characteristics.

To simultaneously motor the motility and ATP levels in the persister population, we sequentially captured brightfield and fluorescent images continuously, as shown in **Movie S7**. Upon analyzing the motility (motile/non-motile) and fluorescence (fluorescent/non-fluorescent) characteristics of 982 persister cells, we identified four distinct states with varied frequency: (1) non-motile but fluorescent (12.23%), (2) motile and fluorescent (10.87%), (3) non-motile and non-fluorescent (24.35%), and (4) motile but non-fluorescent (52.55%). The frequency distribution of these four states is illustrated in **Fig. 6D**. The total percentage of non-fluorescent cells was 76.80%, matching our flow cytometry measurement (low-intensity subpopulation approximately accounted for 73.3% of the whole population, as shown in **fig. S8**). Meanwhile, motile cells accounted for 63.42% of the total persisters, aligning with our previous quantification of motile percentage (74.86%±4.20%) in general. The discrepancies might stem from the significantly lower FPS during the brightfield-fluorescent continuous imaging (approximately 0.35 FPS). The coexistence of four states indicated that ATP levels and cell motility are independent of each other, with no significant correlation. Notably, motile, non-fluorescence cells dominated the entire persister population, further confirming the absence of a positive correlation between motility and ATP. Therefore, these results denied the possibility of ceased movement as a consequence of impaired intracellular ATP utilization.

In sum, rifampin-induced persisters can maintain vigorous glucose uptake levels for a long time. Moreover, they can possess high levels of intracellular ATP, which actually had no significant correlation with motility. Nonetheless, persister cells stopped moving over time, suggesting that the motility was unsustainable by nature and independent of global bioenergetic landscapes.

## Discussion

Exposing mid-exponential phase *E. coli* to bacteriostatic antibiotic rifampin for a brief duration significantly enhances persistence against a lethal dose of ampicillin. While this process involves diverse physiological changes, bacterial motility has remained a relatively underexplored aspect. In this study, we present a comprehensive quantification of cell motility during the formation of rifampin-induced persistence across multiple experimental conditions. Our results revealed that a dynamic remodeling of motility accompanies the transition to persister state, characterized by a reduction in motile percentage, with a decrease in swimming speed and an increase in directional change rate among the motile persister cells.

The observed remodeling of motility during persister formation is not merely a coincidental behavior but actively contributes to the establishment of persistence, as evidenced by the diminished capacity for persister formation upon compromising motility through both the chemical inhibitor CCCP and the physical viscosity agent Ficoll. Mechanistically, we have elucidated that this effect is mediated through the modulation of bactericidal antibiotic uptake, wherein compromised motility during persister formation correlated with elevated intracellular antibiotic accumulation, thus reducing bacterial survival. The direct correlation underscores the functional importance of motility in the persister formation process.

Another central finding of our study is the elucidation of what happens after this initial, motility-dependent stage. We reveal a mechanistic switch from this dynamic defense to a stable, genetically-encoded structural defense state. The transcriptional profile of established persisters demonstrated the dominant downregulation of primary OMP genes data-wise and function-wise. This change effectively “locks the gates” of persister cells, representing a more permanent survival strategy. Therefore, we speculate that the initial motility-based dynamic defense during persister formation buys time for the time-consuming but robust transcriptional defense to activate. The completion of this mechanism switching is functionally confirmed by our observation that motility is no longer essential for persister survival: when motility was abolished in the established persister cells, they remained tolerant to the antibiotic. Furthermore, prior research has indicated that the upregulation of efflux-associated genes played a significant role in persistence, but the OMP genes was not downregulated to collaborate and limit intracellular antibiotic concentrations.^63^ Our findings help complete the picture by showing the critical involvement of OMP genes in persistence and imply potential cooperation between OMP genes and efflux genes might exist.

The dispensability of motility in persistence provided crucial context for its long-term fate. Our prolonged observation revealed that persister motility is not sustained indefinitely; it gradually fades without affecting persister survival. This is consistent with a sustainable strategy for a trait that is no longer required for immediate survival. We attempted to gain insights into the nature of the transient persister motility and its associated physiology. Our findings reveal that persister motility is inherently impermanent, despite the cells maintaining a vigorous ability for nutrient consumption and partially possessing high levels of ATP. Specifically, within the persister population, the ATP levels exhibit bimodal heterogeneity: one subpopulation parallels rifampin-treated cells (higher than mid-exponential phase cells), while the other resembles ampicillin-treated cells (lower than mid-exponential phase cells). The ATP distribution in rifampin-induced persisters against ampicillin thus reflected characteristics of cells treated separately with rifampicin and ampicillin, suggesting a potential additive effect of sequential antibiotic exposure on cellular energy state. Surprisingly, however, correlation analysis between motility and intracellular ATP levels identifies four distinct states, indicating an independence between these two parameters in persisters. Therefore, our results demonstrate that persister motility operates independent of global bioenergetic landscapes, despite the energy demands of flagellar rotation, indicating there might be an involvement of specific, localized energy allocation or diverse mechanisms to power this motility for a limited duration.

Finally, during the last stage of the persister lifecycle, the delayed revival observed upon disrupting motility with both CCCP and Ficoll strongly indicates that motility serves a final, critical purpose. The physical act of movement is crucial for a timely transition back to active growth, likely by facilitating cells to escape from the antibiotic-laden environment and seek out nutrient sources in fresh medium.

In conclusion, our work provides compelling evidence that motility is not merely a characteristic of actively growing cells but follows a crucial, stage-specific lifecycle arc in rifampin-induced persistence: it is fundamental for the entry into the persister state, becomes unnecessary following the establishment of a genetically-encoded structural defense, and is eventually required for an efficient exit from persistence. These findings offer novel insights into the intricate interplay among motility, antibiotic uptake, transcriptional variations, bioenergetic landscapes, and bacterial persistence, opening up new avenues for mechanistic studies. A key unresolved question is the extent to which this motility-dependence represents a generalizable feature of persistence in response to other environmental stressors. However, as antibiotic-cocktail therapies are central to clinical practice, defining the contribution of common cellular processes, such as motility, to antibiotic-induced persistence holds significant translational potential. This perspective suggests, for example, that targeting motility-related processes to eradicate persisters and control chronic bacterial infections may offer a more tractable and effective clinical route than intervening with the complex gene regulatory pathways indicated by previous mechanistic studies. Furthermore, given the functional similarities between bacterial and cancer persister cells (cancer stem cells),^71^ our findings suggest potential strategies to prevent cancer relapse and shortening treatment duration.

## Materials and Methods

### Bacterial Strain, Plasmids and Culture Conditions

The wild-type cells used in this study were *E. coli* K-12 MG1655, purchased from the *E. coli* Genetic Stock Center (Yale University, New Haven).^72^ The ATP reporter plasmid, PPB008 (carrying the rrnBP1-superfolder-GFP-ssrA insert), was constructed previously in our laboratory.

Bacteria were cultured by inoculating single colonies from agar plates into Luria-Bertani broth (LB, containing 1% w/v tryptone, 0.5% w/v yeast extract, and 0.5% w/v NaCl). Cultures were grown at 37°C overnight (approximately 18 hours). Overnight cultures were diluted 1:100 into fresh LB and incubated at 37°C for approximately 2 hours, until they reached mid-exponential phase (optical density at 600 nm ≈ 0.80), prior to experimentation.

### Antibiotics and Chemical Agents

Rifampin (Cas No. 13292-46-1), ampicillin (Cas No. 69-52-3), and CCCP (Cas No. <u>555-60-2</u>) were purchased from Macklin (Catalog No. R6056, A800429, and <u>C838522</u>, respectively).

Ficoll® 400 (Cas No. 26873-85-8) was purchased from Sigma (F8016, lyophilized powder, Type 400-DL, BioReagent, suitable for cell culture). For motility and survival rate assays, Ficoll was added directly to mid-exponential phase bacterial cultures to achieve final concentrations of 5%, 10%, or 15% w/v. To minimize sedimentation of this viscous agent, bacterial cultures were maintained in an incubator shaker until cell suspensions were extracted for experiments.

The potentiometric probe DiOC_2_(3) (Cas No. 905-96-4) was purchased from Aldrich Chemistry (Catalog No. 320684). The fluorescent glucose analog 2-NBDG (Cas No. 186689-07-6) was purchased from TargetMol (Catalog No. T14017).

### Persister Enrichment

Rifampin-induced persisters were generated by exposing mid-exponential phase bacterial cultures to 100 μg/mL rifampin for 30 minutes, followed by the addition of 300 μg/mL ampicillin for overnight incubation (approximately 18 hours).^14^ Given that washing cells to remove rifampin does not affect the results,^23^ and may introduce variability in cell number and motility, this step was omitted prior to ampicillin treatment.

### Survival Rate Determination

To quantify persister cell survival under different experimental conditions, bacterial cultures were serially diluted in LB (10^-2^, 10^-3^, 10^-4^), and 10 μL drops of each dilution were plated onto LB agar. The drops were spread using glass beads. Following overnight incubation at 37°C (approximately 18 hours), images of plates were captured using an iPhone 14 Pro, and colonies were counted using the Jingdian Camera developed by Sunchangsong. Survival rates were calculated by dividing the number of surviving colony-forming units (CFU/mL) by the CFU/mL of the mid-exponential phase culture over six replicates.

### Microscopy Settings

A Nikon Eclipse Ti inverted microscope, equipped with a Plan Apo TIRF 100x Oil DIC H N2 objective and an Andor DU 897 CCD camera, was used for imaging. The objective’s working distance was 0.12 mm (0.16-0.10) at 23°C.

Cell motility imaging was performed at room temperature, maintained by a central air-conditioning system. Continuous time-lapse image acquisition was used to generate cell movement videos. Brightfield image acquisition settings were as follows: exposure time, 10 ms; multiplier, 300; readout speed, 10 MHz; and conversion gain, 1x. Images were acquired at a resolution of 512 x 512 pixels (mono 14-bit), with a pixel resolution of 6.25 pixels per micron. Fluorescent image acquisition settings were: emission wavelength, 561.0 nm; exposure time, 10 ms; multiplier, 30; readout speed, 10 MHz; and conversion gain, 1x.

Real-time image capture settings were identical to those described above, with images taken at 30-minute intervals for selected positions. LB medium containing ampicillin was flowed into the FCS2 chamber using a syringe pump at a speed of 0.85 mL/h.

### Swimming Motion Analysis

Cell motility in liquid environments was recorded using a Bioptechs FCS2 chamber with a 0.2 mm plastic spacer between the slide and coverslip. 8 μL of bacterial suspension was placed into the chamber. Motility was examined for mid-exponential phase wild-type cells, mid-exponential phase cells in LB with 5%, 10%, and 15% w/v Ficoll, mid-exponential phase cells treated with 20 μM CCCP for approximately 2 hours, mid-exponential phase cells after 30 minutes of rifampin pretreatment, rifampin-induced persisters, and persisters treated with CCCP. Videos of swimming cells, at least 15 seconds in duration, were recorded at approximately 22 FPS from three independent experiments and subsequently analyzed.

Motion analysis was proceeded with the TrackMate plugin of Fiji.^43–45^ TrackMate identifies cells across video frames and links them to create tracks. Cell identification was performed using the Cellpose detector.^41,42^ A Cellpose model was trained using 10 bright-field images of mid-exponential phase *E. coli* cells. This model enabled efficient cell identification. In TrackMate, the cell diameter for Cellpose detection was set to 1.8 μm, and no filters were applied to the cell segmentation results. The LAP tracker, suitable for Brownian motion, was used to link cells between adjacent frames, with the following settings: maximum frame-to-frame linking distance, 3.0 μm; maximum track segment gap closing distance, 1.0 μm; and maximum frame gap, 2 frames.

We applied two filters on the tracks from the LAP tracker: track displacement and track minimum speed, to exclude cells adhered to the glass surface and unable to swim freely. The filter threshold was determined by adjusting and examining the trajectories. The threshold values set for mid-exponential phase cells and rifampin-treated cells were: track displacement > 3.00 microns, track minimum speed > 1.01 micron/s. The threshold values set for mid-exponential phase cells in LB with 5%, 10%, and 15% w/v Ficoll were: track displacement > 3.00 microns, track minimum speed > 0.20. The threshold values set for persisters were: track displacement > 3.00, track minimum speed > 0.15. We extracted motility features using the feature analyzers in trackmate. Statistical analysis (distribution plots, mean and standard deviation calculation, ect.) was done using MATLAB and Origin.

### Flow Cytometry Assay

Flow cytometry was performed using an Invitrogen Attune NxT Acoustic Focusing Cytometer. Cells were centrifuged, resuspended in PBS, and measured. Data were modally normalized and results and figures were generated using FlowJo_v10.8.1. Histogram gates were set based on SSC-A vs. FSC-A and FSC-W vs. FSC-H distributions.

### RNA Sequencing

Total RNA sequencing data were first subjected to quality assessment using FastQC, followed by trimming of adapter sequences and low-quality reads using Trimmomatic, ensuring the retention of high-quality, clean reads for downstream analysis. Clean reads were then aligned to the E. coli reference genome using Bowtie2, and alignment statistics were compiled to assess mapping performance. Additional quality metrics, including redundancy analysis and insert size distribution, were evaluated with RSeQC, while Qualimap was used to assess coverage uniformity and distribution across genomic regions. Gene coverage and genome-wide read distribution were further analyzed using BEDTools. Subsequently, gene expression levels were quantified using featureCounts based on existing gene models. Differential gene expression was assessed using DESeq2. Functional enrichment of differentially expressed genes was performed using topGO for Gene Ontology (GO) terms, while clusterProfiler was used for KEGG pathway and COG category enrichment.

## Notes

### Note 1

As we examined the motility features of cell populations across different conditions -- mid-exponential phase cells, rifampin treated cells, persisters, mid-exponential phase cells in LB with 5%, 10%, and 15% w/v Ficoll) -- we consistently observed a negative correlation between swimming speed and directional change rate. As shown in **Fig. 2E**, **fig. S2** and **S5**, in each assessed population, cells with slower swimming speeds exhibited higher directional change rates. This negative correlation prompts consideration of whether the observed increase in directional change rate during persister formation was solely attributable to the reduction in speed.

Previous research has indicated that near planar glass surfaces in unstimulated environment, *E. coli* cells typically swim in circular trajectories.^29^ Given our experiment setup, including the working distance of the objective and the chamber coverslip width, then the distance between the observed cells and the glass surface should be further than 10 μm but less than 400 μm. Under these conditions, the cells could not be considered to swim completely free of surface effects in a liquid phase and the bacteria-surface interactions influenced their movement.^73^ Faster-swimming *E. coli* cells generally swim in circular trajectories with larger diameters,^74^ while slower-swimming cells show the opposite trend. Therefore, the negative correlation between speed and directional change rate observed in our results could be interpreted within this framework of surface-influenced motility. The relationship between the mean speed and mean directional change across all measured populations is shown in **fig. S9**. While the data point representing rifampin-treated cells (*R*) aligns with the monotonic negative relationship, the data point representing persisters (*R-A*) appears as an outlier, exhibiting a notable deviation. This implies that the primary alteration in motility following rifampin pretreatment was the decrease in speed. As regards persisters, although our experimental setup likely contributed to some increase in directional change rate with respect to the decrease in speed, our findings indicate that persisters possess an inherent propensity for more frequent changes in swimming direction and more circular trajectory.

### Note 2

To determine whether nutrient depletion contributed to the loss of motility, we transferred the rifampin-induced persisters to fresh LB medium supplemented with the same concentrations of antibiotics after overnight ampicillin exposure and monitored cell behavior over time (shown in **Movie S5**). **fig. S10a** illustrated the cell count and motile percentage of samples from this culture at five distinct time points. The average cell number fluctuated over time with no statistically significant difference (**fig. S10b**), indicating no substantial cell lysis or death. However, the motile percentage dropped from about 70% to nearly 0% (the smaller motile percentage at the beginning might be contributed to the relocation process which involved centrifuge and resuspension). This observation was consistent with the behavior of cells in the original culture, suggesting that nutrient depletion was not the cause for the unsustainable motility in persister cells. Furthermore, we evaluated the influence of secondary metabolites by relocating the persister cells to PBS to inhibit cellular metabolism and subsequently repeating the prolonged observation experiment (**Movie S6**). Interestingly, as illustrated in **fig. S10c**, the temporal dynamics of both cell count and motile percentage were similar to those from the original culture or the fresh culture, with the cell count remaining stable (**fig. S10d**) and motile percentage ultimately dropping to almost 0%. Collectively, these results suggest a common behavioral pattern that persister cells progressively cease motility over time, regardless of external environmental conditions. Moreover, persister cells demonstrate sustained antibiotic tolerance despite the loss of motility.

## Acknowledgments

The graphical abstract, Fig. 1A, and Fig. 4F were created with Figdraw.

## Funding

National Natural Science Foundation of China NO. 32171245 (XF, YX)

National Key Research and Development Program of China 2024YFA 0919600 (XF, YX)

## Author contributions

Conceptualization: XF, JW

Methodology: XF, YX, JW

Investigation: YX

Visualization: YX

Funding acquisition: XF, JW

Project administration: XF, JW

Supervision: XF, JW

Writing – original draft: YX

Writing – review & editing: YX, XF, JW

## Data and materials availability

The experimental materials and original data are available from the corresponding authors upon reasonable request.

**Supplementary Materials** Supplementary Figures S1 to S10 Movies S1 to S7

