## Supplementary Materials for "Motility Functions as an Essential Defense in the Bacterial Persister Lifecycle"


Yixiao Xiong *et al.*

*Xiaona Fang.
*Jin Wang.


**This PDF file includes:**

figs. S1 to S10

**Other Supplementary Materials for this manuscript include the following:**

Movies S1 to S7

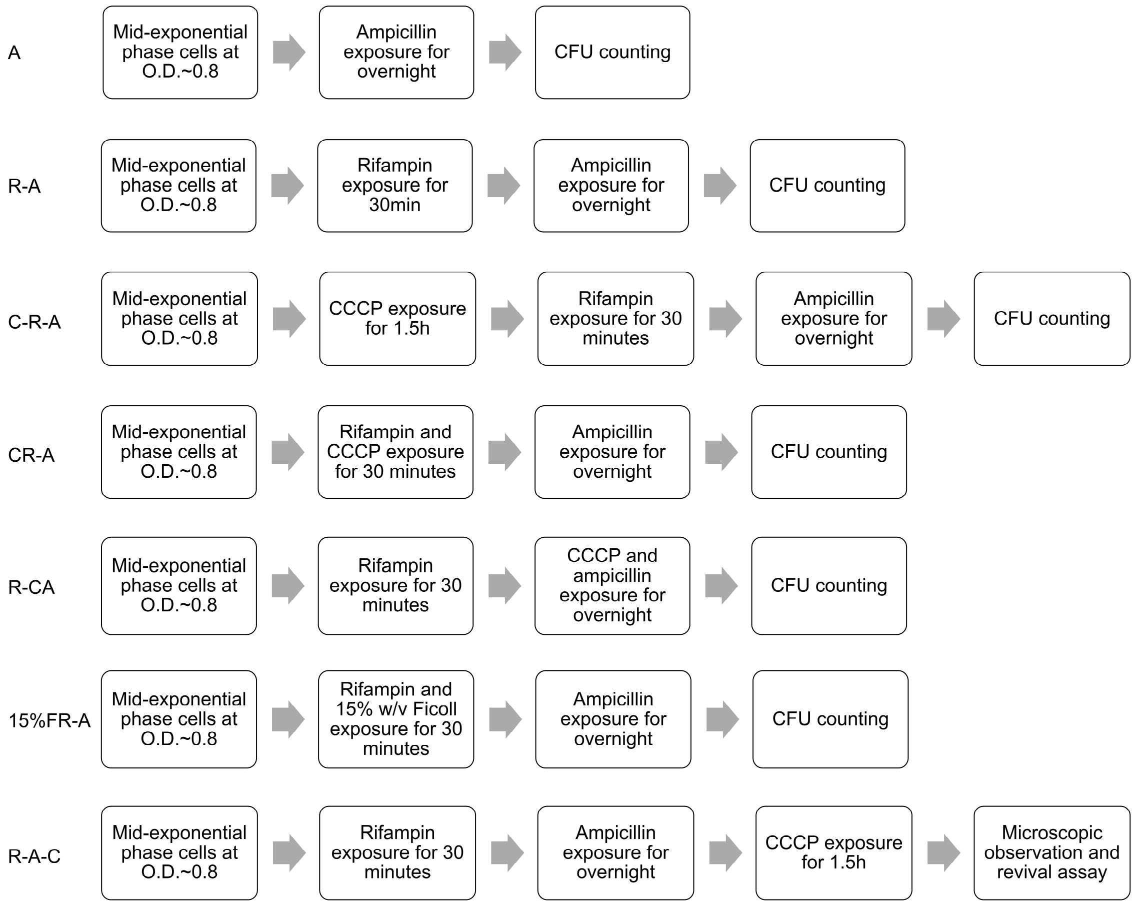


**fig. S1.** Schematic of protocols used for persister generation and motility modification.

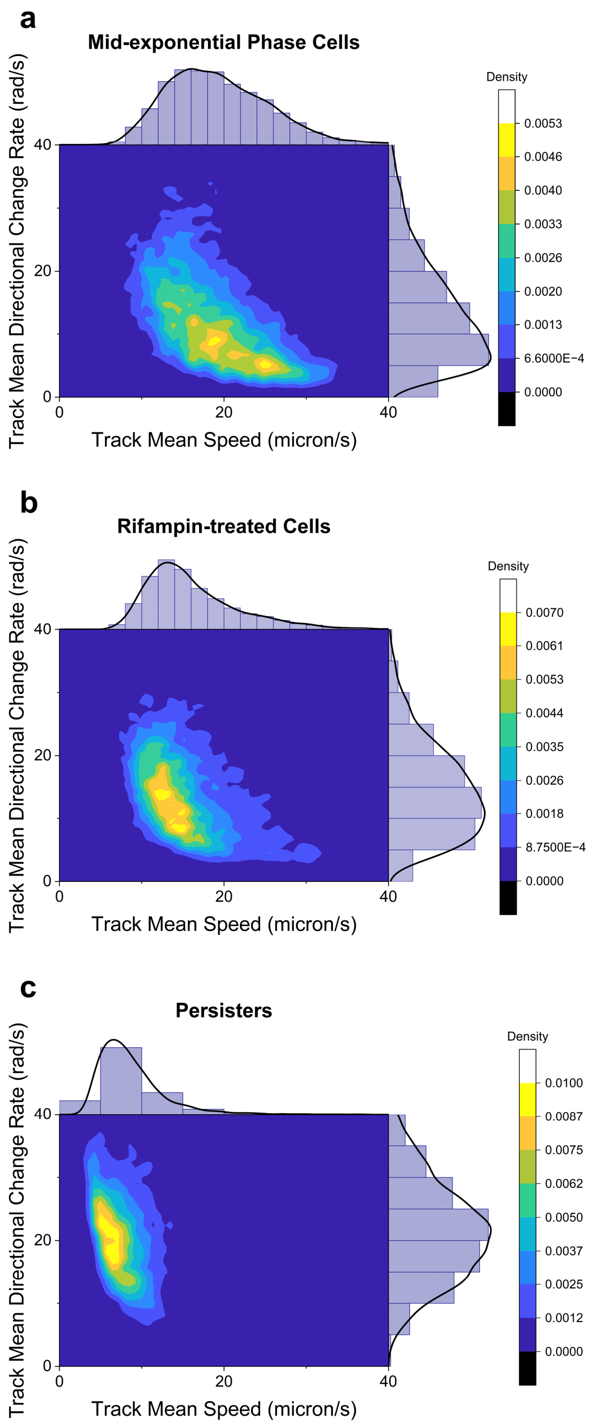


**fig. S2.** Two-dimensional distributions correlating track mean speed and mean directional change rate for mid-exponential phase cells (a), rifampin-treated cells (b), and rifampin-induced persisters (c), with marginal distribution plots.

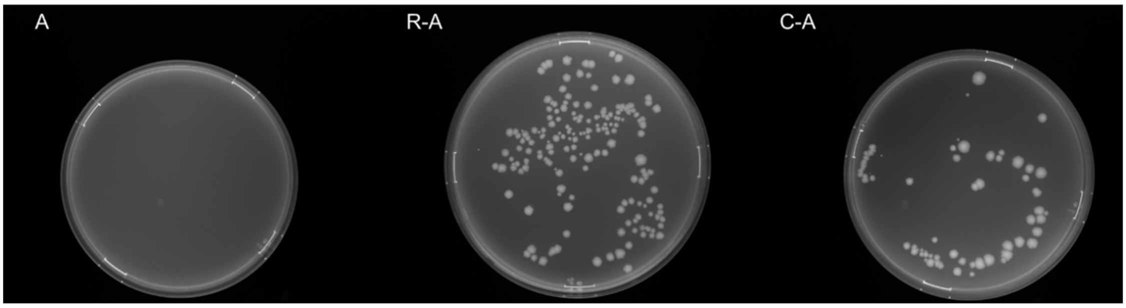


**fig. S3.** Representative results of survival rate against ampicillin without pretreatment (*A*, left plate), with rifampin pretreatment (*R-A*, middle plate), and with CCCP pretreatment (*C-A*, right plate). The samples were drawn from the same culture before pretreatment and ampicillin treatment. The plates were incubated at 37°C overnight after plating with 10 μL drops of 1:100 LB diluted culture.

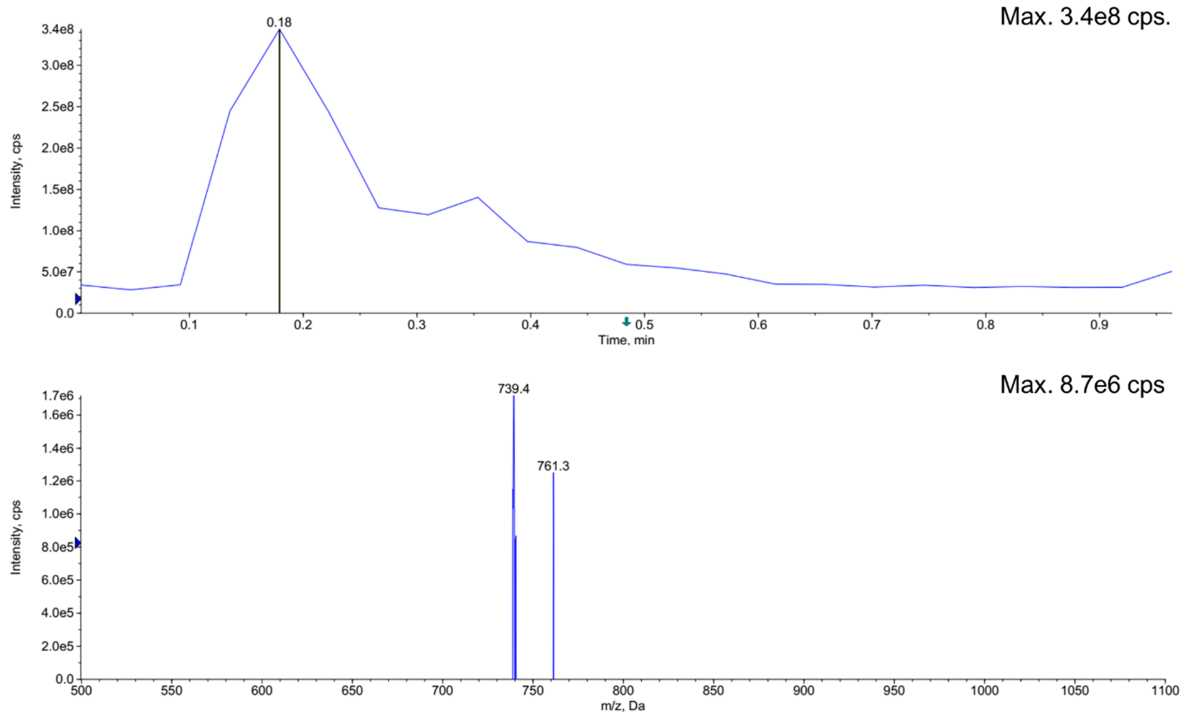


**fig. S4.** Mass spectrometry of FITC-ampicillin.

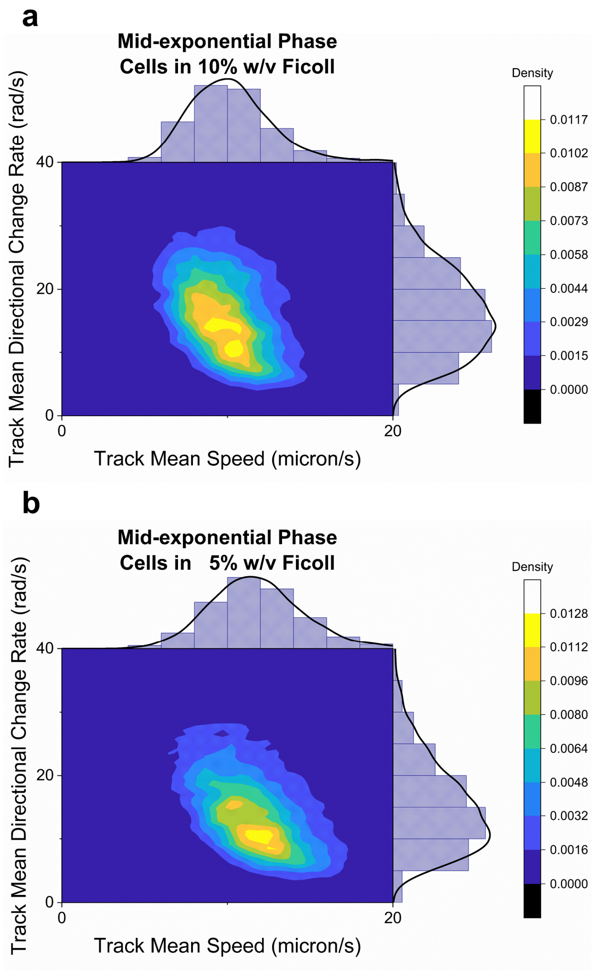


**fig. S5.** Two-dimensional distributions correlating track mean speed and mean directional change rate for mid-exponential phase cells swimming in LB supplemented with 10% (a) and 5% w/v Ficoll (b), with marginal distribution plots.

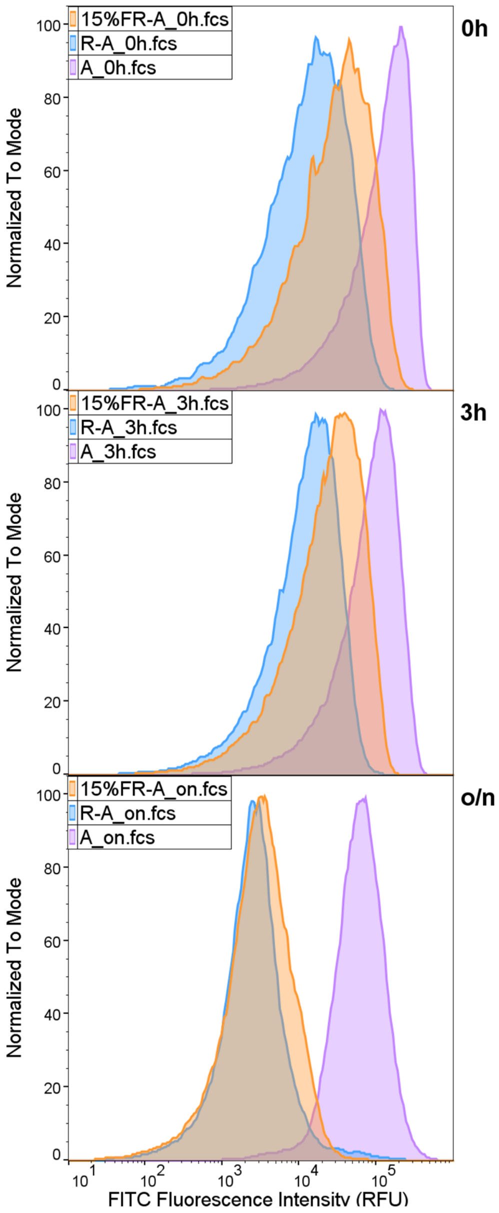


**fig. S6.** Intracellular antibiotic uptake of FITC-ampicillin at the onset (0h, top panel), early phase (3h, middle panel), and the conclusion (o/n, bottom panel) of ampicillin exposure, in rifampin-induced persisters with intact motility during rifampin-pretreatment (*R-A,* blue distribution), rififampin-induced persisters under the influence of 15% w/v Ficoll during rifampin-pretreatment (15%F*R-A,* orange distribution), and cells directly exposed to ampicillin (*A,* purple distribution), respectively. The frequency distributions are all normalized to mode.

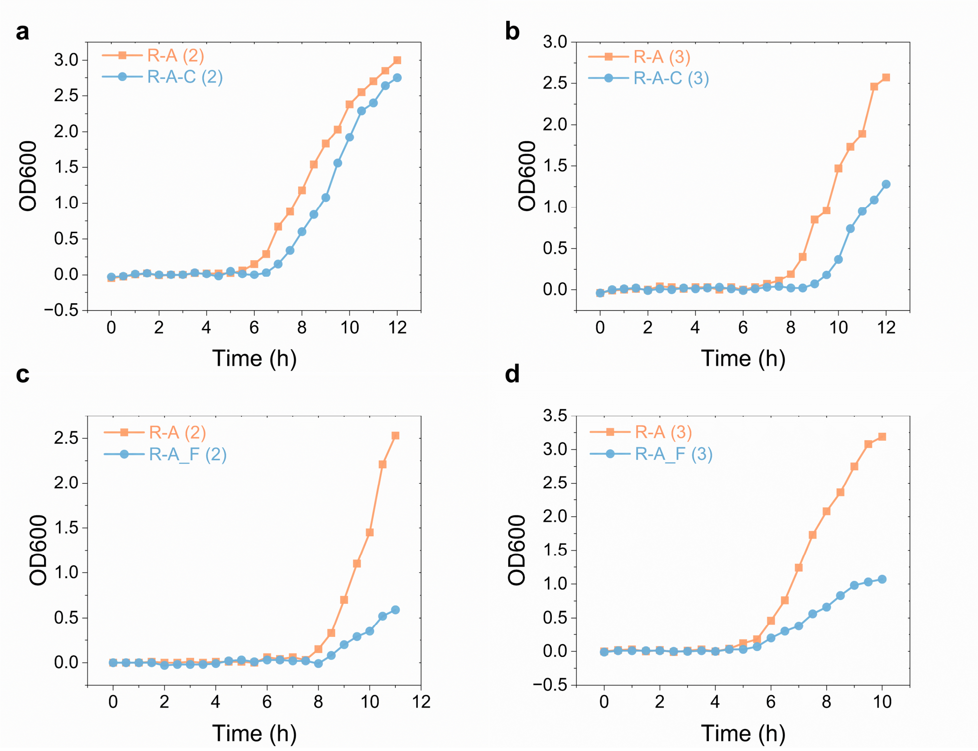


**fig. S7.** Representative growth curves of persister cells (*R-A*) and persister cells exposed to CCCP (*R-A-C*), or persister cells resuming growth in fresh medium with (*R-A_F*) or without (*R-A*) 15% w/v Ficoll, during revival in fresh medium. The curves in each plot represent results from an independent experiment.

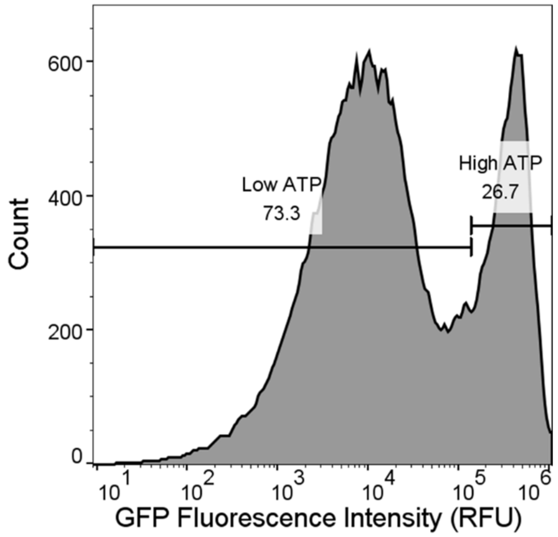


**fig. S8.** Percentages of the two subpopulations exhibiting high and low ATP, indicated by GFP fluorescence measured via flow cytometry. The numbers on the subpopulations represent percentages.

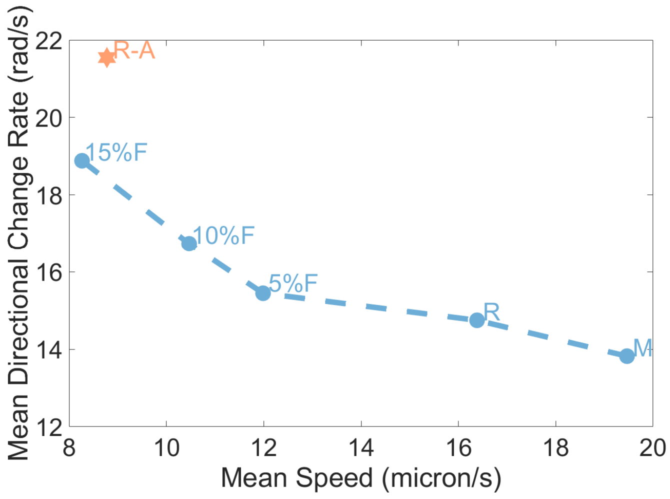


**fig. S9.** The correlation between mean directional change rate and mean speed of all the cell populations examined in this study. Each data point represents a cell population in a particular state or under a particular experimental condition. The dashed line indicates the potential influence of the experimental setup on bacterial motility behaviors. The data representing persister cells is aberrant compared to other data.

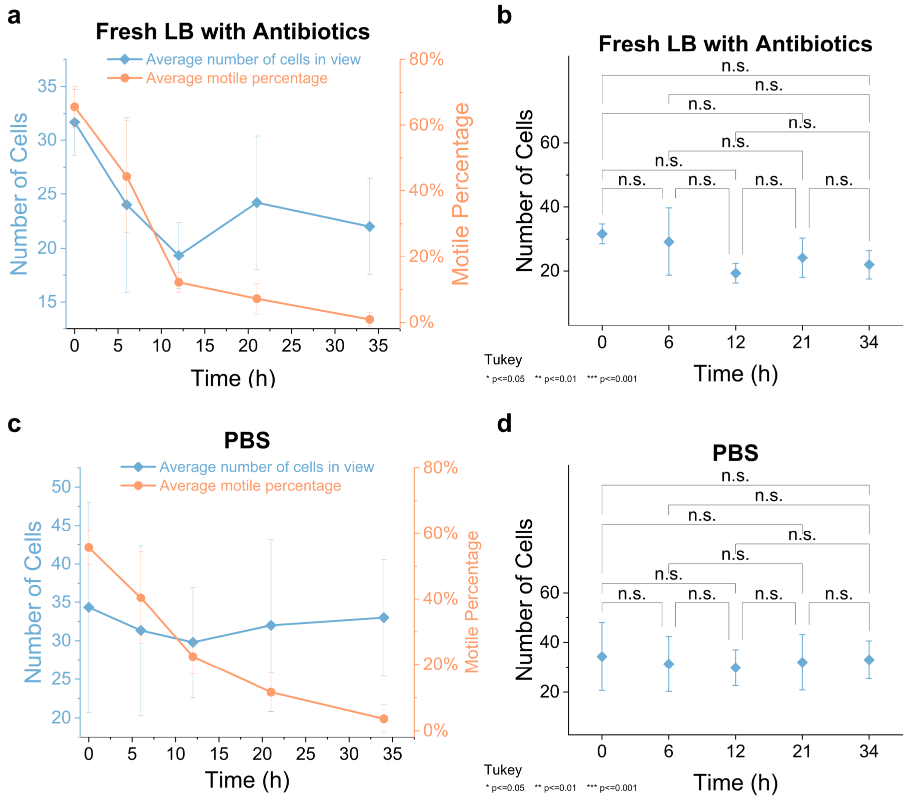


**fig. S10.** The cell counts and motile percentage of persister cells under different external culture conditions during prolonged observation. **a)** Cell count (blue, left axis) and motile percentage (orange, right axis) of persisters in fresh LB with corresponding antibiotics at 0h, 6h, 12h, 21h, 34h after relocation from original culture (approximately 16h, 22h, 28h, 37h, 50h after rifampin pretreatment). Data points represent the mean (±standard deviation) from at least three fields of view. **b)** Tukey paired comparison results of number of persister cells in fresh LB with antibiotics after 0h, 6h, 12h, 21h, 34h. At the 0.05 level, Shapiro-Wilk normality test suggests that the data at each time point was significantly drawn from a normally distributed population. The abbreviation 'n.s.' indicates no statistically significant difference between the samples in a test (all P values > 0.05). **c**) Cell count (blue, left axis) and motile percentage (orange, right axis) of persisters in PBS at 0h, 6h, 12h, 21h, 34h after relocation from original culture (approximately 16h, 22h, 28h, 37h, 50h after rifampin pretreatment). Data points represent the mean (±standard deviation) from at least three fields of view. **d)** Tukey paired comparison results of number of persister cells in PBS after 0h, 6h, 12h, 21h, 34h. At the 0.05 level, Shapiro-Wilk normality test suggests that the data at each time point was significantly drawn from a normally distributed population. The abbreviation 'n.s.' indicates no statistically significant difference between the samples in a test (all P values > 0.05).

**Movie S1 (separate file).** Motility of mid-exponential phase cells (a), rifampin-treated cells (b), rifampin-induced persisters (c), and ampicillin-treated cells (d) in their original culture environment, observed in FCS2 chamber.

**Movie S2 (separate file).** Motility of CCCP-treated cells (a), cells in LB with 15% (b), 10% (c), and 5% (d) w/v Ficoll, observed in FCS2 chamber.

**Movie S3 (separate file).** Motility of rifampin-induced persister following exposure to CCCP, observed in FCS2 chamber.

**Movie S4 (separate file).** Persister cells in their original culture after 0h, 6h, 12h, 21h, and 34h.

**Movie S5 (separate file).** Persister cells in fresh LB with antibiotics after 0h, 6h, 12h, 21h, and 34h.

**Movie S6 (separate file).** Persister cells in PBS after 0h, 6h, 12h, 21h, and 34h.

**Movie S7 (separate file).** Overlay video of consecutive bright-field and fluorescence microscopy imaging at 488nm for persister cells with fluorescence ATP reporter.
